## Supplementary material for "Analytical solution of linearized equations of the Morris-Lecar neuron model at large constant stimulation": Two supplementary Figures, MATLAB scripts, and generated data used for Figures: ML_neuron_dynamics_ode23s.m

```
clear all;

% potential V is in mV
% time t is in ms
% electical capacity C_m is in µF/cm^2
% conductivity g is in mS/cm^2
% stimulating current I_stim is in µA/cm^2

% Parameters for the Morris-Lecar model were taken from J. Rinzel, B. Ermentrout, Analysis of neural excitability and oscillations. 
% In: C. Koch, I. Segev (Eds.), Methods in neuronal modeling: From synapses to networks (2nd Ed., MIT Press, Cambridge, 1998), pp. 251-291.

C_m = 20; % µF/cm^2

g_K = 8; % mS/cm^2
g_L = 2; % mS/cm^2

V_Ca = 120; % mV
V_K = -84; % mV
V_L = -60; % mV
V1 = -1.2; % mV
V2 = 18; % mV

excitability_type = 1;

if (excitability_type == 1)
    g_Ca = 4; % mS/cm^2
    V3 = 12; % mV
    V4 = 17.4; % mV
    beta = 0.067; % kHz, beta = 1/tau_max, tau_max = 14.925 ms
end;

if (excitability_type == 2)
    g_Ca = 4.4; % mS/cm^2
    V3 = 2; % mV
    V4 = 30; % mV
    beta = 0.04; % kHz, beta = 1/tau_max, tau_max = 25 ms
end;

%-------------- Functions------------------

m_inf = @(V) 0.5.*(1 + tanh((V-V1)./V2)); 
w_inf = @(V) 0.5.*(1 + tanh((V-V3)./V4)); 
tau_w = @(V) 1./cosh((V-V3)./(2.*V4)); 

rest = @(V) (g_Ca*m_inf(V)*(-V + V_Ca) + g_K*w_inf(V)*(-V + V_K) + g_L*(-V + V_L))./C_m;

%---------- Finding V_rest  ----------- %
V_start = -60; % mV 
format long
V_rest = fzero(rest,V_start) % sum of ion currents = 0
rest(V_rest)
%--------------------------%

t_max = 550; % ms;
I_stim = 118; % µA/cm^2 

% Below V = y(1) and w = y(2):

F = @(t,y) [(g_Ca.*m_inf(y(1)).*(-y(1) + V_Ca) + g_K.*y(2).*(-y(1) + V_K) + g_L.*(-y(1) + V_L) + I_stim)./C_m; beta.*(w_inf(y(1))-y(2))./tau_w(y(1))];

[T, Y] = ode23s(F,[0 t_max],[V_rest w_inf(V_rest)]);

figure(1)
plot(T,Y(:,1),'blue')
xlabel('t [ms]')
ylabel('V [mV]')

figure(2)
plot(T,Y(:,2),'red')
xlabel('t [ms]')
ylabel('w')

fid = fopen('ML_neuron_V(t)_ode23s_ExcType=1_I_stim=118.txt','wt');
OUT = [T, Y(:,1)];
fprintf(fid,'%10.12f %10.12f\n',OUT');
fclose(fid);
```
