## Supplementary material for "Analytical solution of linearized equations of the Morris-Lecar neuron model at large constant stimulation": Two supplementary Figures, MATLAB scripts, and generated data used for Figures: ML_neuron_dynamics_ode23s_FITTING_analyt_solution.m

```
clear all;

% potential V is in mV
% time t is in ms
% electical capacity C_m is in µF/cm^2
% conductivity g is in mS/cm^2
% stimulating current I_stim is in µA/cm^2

% Parameters for the Morris-Lecar model were taken from J. Rinzel, B. Ermentrout, Analysis of neural excitability and oscillations. 
% In: C. Koch, I. Segev (Eds.), Methods in neuronal modeling: From synapses to networks (2nd Ed., MIT Press, Cambridge, 1998), pp. 251-291.

C_m = 20; % µF/cm^2

g_K = 8; % mS/cm^2
g_L = 2; % mS/cm^2

V_Ca = 120; % mV
V_K = -84; % mV
V_L = -60; % mV
V1 = -1.2; % mV
V2 = 18; % mV

excitability_type = 1;

if (excitability_type == 1)
    g_Ca = 4; % mS/cm^2
    V3 = 12; % mV
    V4 = 17.4; % mV
    beta = 0.067; % kHz, beta = 1/tau_max, tau_max = 14.925 ms
end;

if (excitability_type == 2)
    g_Ca = 4.4; % mS/cm^2
    V3 = 2; % mV
    V4 = 30; % mV
    beta = 0.04; % kHz, beta = 1/tau_max, tau_max = 25 ms
end;

%-------------- Functions------------------

m_inf = @(V) 0.5.*(1 + tanh((V-V1)./V2)); 
w_inf = @(V) 0.5.*(1 + tanh((V-V3)./V4)); 
tau_w = @(V) 1./cosh((V-V3)./(2.*V4)); 

rest = @(V) (g_Ca*m_inf(V)*(-V + V_Ca) + g_K*w_inf(V)*(-V + V_K) + g_L*(-V + V_L));

%---------- Finding V_rest  ----------- %
V_start = -60; % mV 
format long
V_rest = fzero(rest,V_start) % sum of ion currents = 0
rest(V_rest)
%--------------------------%

t_max = 1200; % ms;
I_stim = 116.3; % µA/cm^2 

% Below V = y(1) and w = y(2):

F = @(t,y) [(g_Ca.*m_inf(y(1)).*(-y(1) + V_Ca) + g_K.*y(2).*(-y(1) + V_K) + g_L.*(-y(1) + V_L) + I_stim)./C_m; beta.*(w_inf(y(1))-y(2))./tau_w(y(1))];

[T, Y] = ode23s(F,[0 t_max],[V_rest w_inf(V_rest)]);

V = Y(:,1);
w = Y(:,2);

fid = fopen('ML_neuron_V(t)_ode23s_ExcType=1_I_stim=116.3.txt','wt');
OUT = [T, V];
fprintf(fid,'%10.12f %10.12f\n',OUT');
fclose(fid); 

fid = fopen('ML_neuron_w(t)_ode23s_ExcType=1_I_stim=116.3.txt','wt');
OUT = [T, w];
fprintf(fid,'%10.12f %10.12f\n',OUT');
fclose(fid); 

%============ Comparison with analytical solution =========================

stat = @(V) (g_Ca*m_inf(V)*(-V + V_Ca) + g_K*w_inf(V)*(-V + V_K) + g_L*(-V + V_L) + I_stim);

V_start = 10; % mV 
format long
V_st = fzero(stat,V_start) % steady solution of equation dV/dt = 0
stat(V_st)

a = w_inf(V_st)

b = (0.5./V4).*1/(cosh((V_st-V3)/V4))^2 % (mV)^(-1)

p = m_inf(V_st)

q = (0.5./V2).*1/(cosh((V_st-V1)/V2))^2 % (mV)^(-1)

A = (g_Ca*(p + q*(V_st - V_Ca)) + g_K*a + g_L)/C_m % kHz = 10^3 S/F

B = g_K*b*(V_st - V_K)/C_m % kHz

tau = tau_w(V_st)/beta % ms

%============================%

% V(t = 693.3 ms) = 16.35 mV <- target maximum for I_stim = 116.3 µA/cm^2

t_0 = 693.3; % ms

V_0 = 16.35; % mV

U_0 = V_0 - V_st

w_0 = (I_stim - g_Ca*m_inf(V_0)*(V_0 - V_Ca) - g_L*(V_0 - V_L))./(g_K*(V_0 - V_K))

A_K = g_K*(w_0 - a)/C_m

Y = A + 1/tau % kHz

Z = (A + B)/tau % (kHz)^2

gamma = Y/2 % kHz

omega_0 = sqrt(Z) % kHz

omega = sqrt((omega_0)^2 - gamma^2) % kHz

dt = 0.1 % ms

t = t_0:dt:t_max;

U = U_0.*exp(-gamma.*(t-t_0)).*(cos(omega.*(t-t_0)) + (gamma/omega).*sin(omega.*(t-t_0)));

ita = atan(gamma./omega)

U1 = U_0.*exp(-gamma.*(t-t_0)).*cos(omega.*(t-t_0) - ita)./cos(ita);

fid = fopen('ML_neuron_approx_V(t)_ExcType=1_I_stim=116.3.txt','wt');
OUT = [t; (U+V_st)];
fprintf(fid,'%10.12f %10.12f\n',OUT);
fclose(fid);  

figure(1)
plot(T,V,'blue',t,U+V_st,'green',t,U1+V_st,'red')
xlabel('t [ms]')
ylabel('V [mV]')

w_1 = b.*U_0.*(A./B);

H = ((omega_0)^2)./(omega.*A) - gamma./omega

w_theor = a + (w_0 - a + w_1).*exp(-(t-t_0)./tau) + w_1.*exp(-gamma.*(t-t_0)).*(-cos(omega.*(t-t_0)) + H.*sin(omega.*(t-t_0)));

xi = atan((1 - gamma.*tau)./(omega.*tau))

W_a = (b.*U_0./(omega.*tau)).*sqrt(1 + A./B)

W_c = w_0 - a - W_a.*sin(xi-ita)

w_theor2 = a + W_c.*exp(-(t-t_0)./tau) + W_a.*exp(-gamma.*(t-t_0)).*sin(omega.*(t-t_0) + xi - ita);

fid = fopen('ML_neuron_approx_w(t)_ExcType=1_I_stim=116.3.txt','wt');
OUT = [t; w_theor];
fprintf(fid,'%10.12f %10.12f\n',OUT);
fclose(fid); 

figure(2)
plot(T,w,'black',t,w_theor,'green',t,w_theor2,'red')
xlabel('t [ms]')
ylabel('w')
```
