## Supplementary material for "Analytical solution of linearized equations of the Morris-Lecar neuron model at large constant stimulation": Two supplementary Figures, MATLAB scripts, and generated data used for Figures: ML_neuron_dynamics_ode23s_FITTING_goodness_of_fit.m

```
clear all;

% potential V is in mV
% time t is in ms
% electical capacity C_m is in µF/cm^2
% conductivity g is in mS/cm^2
% stimulating current I_stim is in µA/cm^2

% Parameters for Morris-Lecar were taken from J. Rinzel, B. Ermentrout, Analysis of neural excitability and oscillations. 
% In: C. Koch, I. Segev (Eds.), Methods in neuronal modeling: From synapses to networks (2nd Ed., MIT Press, Cambridge, 1998), pp. 251-291.

C_m = 20; % µF/cm^2

g_K = 8; % mS/cm^2
g_L = 2; % mS/cm^2

V_Ca = 120; % mV
V_K = -84; % mV
V_L = -60; % mV
V1 = -1.2; % mV
V2 = 18; % mV

excitability_type = 1;

if (excitability_type == 1)
    g_Ca = 4; % mS/cm^2
    V3 = 12; % mV
    V4 = 17.4; % mV
    beta = 0.067; % kHz, beta = 1/tau_max, tau_max = 14.925 ms
end;

if (excitability_type == 2)
    g_Ca = 4.4; % mS/cm^2
    V3 = 2; % mV
    V4 = 30; % mV
    beta = 0.04; % kHz, beta = 1/tau_max, tau_max = 25 ms
end;

%-------------- Functions------------------

m_inf = @(V) 0.5.*(1 + tanh((V-V1)./V2)); 
w_inf = @(V) 0.5.*(1 + tanh((V-V3)./V4)); 
tau_w = @(V) 1./cosh((V-V3)./(2.*V4)); 

rest = @(V) (g_Ca*m_inf(V)*(-V + V_Ca) + g_K*w_inf(V)*(-V + V_K) + g_L*(-V + V_L));

%---------- Finding V_rest  ----------- %
V_start = -60; % mV 
format long
V_rest = fzero(rest,V_start) % sum of ion currents = 0
rest(V_rest)
%--------------------------%

t_max = 1000; % ms;
I_stim = 116.3; % µA/cm^2 

% Below V = y(1) and w = y(2):

F = @(t,y) [(g_Ca.*m_inf(y(1)).*(-y(1) + V_Ca) + g_K.*y(2).*(-y(1) + V_K) + g_L.*(-y(1) + V_L) + I_stim)./C_m; beta.*(w_inf(y(1))-y(2))./tau_w(y(1))];

[T, Y] = ode23s(F,[0 t_max],[V_rest w_inf(V_rest)]);

V = Y(:,1);
w = Y(:,2);

%===============================

stat = @(V) (g_Ca*m_inf(V)*(-V + V_Ca) + g_K*w_inf(V)*(-V + V_K) + g_L*(-V + V_L) + I_stim);

V_start = 10; % mV 
format long
V_st = fzero(stat,V_start) % steady solution of equation dV/dt = 0
stat(V_st)

maxV = zeros(1,1);
maxT = zeros(1,1);
errV = 0.001; % mV
p = 0;

for i=2:(length(V)-1)

 if (V(i)>V(i-1))&&(V(i)>V(i+1))&&(abs(V(i)-V(i+1))>errV)&&(abs(V(i)-V_st)>0.1)  
     p = p + 1;
     maxV(p) = V(i);
     maxT(p) = T(i);
 end;                            
   
end;

%============ Comparison with analytical solution =========================

a = w_inf(V_st);

b = (0.5./V4).*1/(cosh((V_st-V3)/V4))^2; % (mV)^(-1)

p = m_inf(V_st);

q = (0.5./V2).*1/(cosh((V_st-V1)/V2))^2; % (mV)^(-1)

A = (g_Ca*(p + q*(V_st - V_Ca)) + g_K*a + g_L)/C_m; % kHz = 10^3 S/F

B = g_K*b*(V_st - V_K)/C_m; % kHz

tau = tau_w(V_st)/beta; % ms

%============================%

S = zeros(1,1);
SS = zeros(1,1);
deltaV_0 = zeros(1,1);
t_0_rec = zeros(1,1);
R_squared = zeros(1,1);

fid = fopen('t0_V0_Vst_deltaV0_S_SS_R2.txt','w');

t_0_min = 500; % ms
t_0_max = 868; %ms

i_good = find((maxT>=t_0_min)&(maxT<=t_0_max));

for i=min(i_good):1:max(i_good)
    
t_0 = maxT(i);
V_0 = maxV(i);    

t = T(find(T>=t_0));
V_cut = V(find(T>=t_0));

U_0 = V_0 - V_st;

Y = A + 1/tau; % kHz

Z = (A + B)/tau; % (kHz)^2

gamma = Y/2; % kHz

omega_0 = sqrt(Z); % kHz

omega = sqrt((omega_0)^2 - gamma^2); % kHz

ita = atan(gamma./omega);

U = U_0.*exp(-gamma.*(t-t_0)).*cos(omega.*(t-t_0) - ita)./cos(ita);

w_0 = (I_stim - g_Ca*m_inf(V_0)*(V_0 - V_Ca) - g_L*(V_0 - V_L))./(g_K*(V_0 - V_K));

xi = atan((1 - gamma.*tau)./(omega.*tau));

W_a = (b.*U_0./(omega.*tau)).*sqrt(1 + A./B);

W_c = w_0 - a - W_a.*sin(xi-ita);

w_theor = a + W_c.*exp(-(t-t_0)./tau) + W_a.*exp(-gamma.*(t-t_0)).*sin(omega.*(t-t_0) + xi - ita);

%======================================

residuals = abs(V_cut - (V_st + U));

j = i - min(i_good) + 1;

S(j) = sum(residuals)/length(residuals);

SS(j) = S(j)/(V_0 - V_st);

deltaV_0(j) = (V_0-V_st)/V_st;

t_0_rec(j) = t_0;

R_squared(j) = sum((V_st + U - mean(V_cut)).^2)/sum((V_cut - mean(V_cut)).^2);

OUT = [t_0, V_0, V_st, deltaV_0(j), S(j), SS(j), R_squared(j)];
fprintf(fid,'%10.12f %10.12f %10.12f %10.12f %10.12f %10.12f %10.12f\n',OUT');

end;

fclose(fid);

figure(1)
plot(t_0_rec,deltaV_0,'-* blue')
xlabel('t_{0} [ms]')
ylabel('(V_{0}-V_{st})/V_{st}')

figure(2)
plot(t_0_rec,S,'-* blue')
xlabel('t_{0} [ms]')
ylabel('S [mV]')

figure(3)
plot(deltaV_0,S,'-* blue')
xlabel('(V_{0}-V_{st})/V_{st}')
ylabel('S [mV]')

figure(4)
plot(t_0_rec,SS,'-* blue')
xlabel('t_{0} [ms]')
ylabel('S/(V_{0}-V_{st})')

figure(5)
plot(deltaV_0,SS,'-* blue')
xlabel('(V_{0}-V_{st})/V_{st}')
ylabel('S/(V_{0}-V_{st})')

figure(6)
plot(t_0_rec,R_squared,'-* blue')
xlabel('t_{0} [ms]')
ylabel('R^{2}')

figure(7)
plot(deltaV_0,R_squared,'-* blue')
xlabel('(V_{0}-V_{st})/V_{st}')
ylabel('R^{2}')
```
