## Supplementary material for "Analytical solution of linearized equations of the Morris-Lecar neuron model at large constant stimulation": Two supplementary Figures, MATLAB scripts, and generated data used for Figures: ML_neuron_damped_lin_eq_parameters_of_I_stim.m

```
clear all;

% potential V is in mV
% time t is in ms
% electical capacity C_m is in µF/cm^2
% conductivity g is in mS/cm^2
% stimulating current I_stim is in µA/cm^2

% Parameters for the Morris-Lecar model were taken from J. Rinzel, B. Ermentrout, Analysis of neural excitability and oscillations. 
% In: C. Koch, I. Segev (Eds.), Methods in neuronal modeling: From synapses to networks (2nd Ed., MIT Press, Cambridge, 1998), pp. 251-291.

C_m = 20; % µF/cm^2

g_K = 8; % mS/cm^2
g_L = 2; % mS/cm^2

V_Ca = 120; % mV
V_K = -84; % mV
V_L = -60; % mV
V1 = -1.2; % mV
V2 = 18; % mV

excitability_type = 1;

if (excitability_type == 1)
    g_Ca = 4; % mS/cm^2
    V3 = 12; % mV
    V4 = 17.4; % mV
    beta = 0.067; % kHz, beta = 1/tau_max, tau_max = 14.925 ms
end;

if (excitability_type == 2)
    g_Ca = 4.4; % mS/cm^2
    V3 = 2; % mV
    V4 = 30; % mV
    beta = 0.04; % kHz, beta = 1/tau_max, tau_max = 25 ms
end;

%-------------- Functions ----------------%

m_inf = @(V) 0.5.*(1 + tanh((V-V1)./V2)); 
w_inf = @(V) 0.5.*(1 + tanh((V-V3)./V4)); 
tau_w = @(V) 1./cosh((V-V3)./(2.*V4)); 

%----------------------------------------%
format long

% I_stim is constant stimulating current, µA/cm^2
I_stim_min = 0;
dI_stim = 0.01;
I_stim_max = 200;

k = 0;
I_stim_rec = zeros(1,1);
V_st_rec = zeros(1,1);
A = zeros(1,1);
B = zeros(1,1);
tau = zeros(1,1);
Y = zeros(1,1);
Z = zeros(1,1); 
omega_0 = zeros(1,1);
gamma = zeros(1,1);
omega = zeros(1,1);
Im_omega = zeros(1,1);

fid = fopen('lin_eq_parameters_of_I_stim_ML_neuron_ExcType=1.txt','wt');

fid2 = fopen('omegas_and_gamma_of_I_stim_ML_neuron_ExcType=1.txt','wt');

fid3 = fopen('negative_part_gamma_of_I_stim_ML_neuron_ExcType=1.txt','wt');

fid4 = fopen('positive_part_gamma_of_I_stim_ML_neuron_ExcType=1.txt','wt');

fid5 = fopen('omega0_imaginary_part_of_I_stim_ML_neuron_ExcType=1.txt','wt');

fid6 = fopen('omega0_of_I_stim_ML_neuron_ExcType=1.txt','wt');

fid7 = fopen('omega_imaginary_part_of_I_stim_ML_neuron_ExcType=1.txt','wt');

fid8 = fopen('omega_of_I_stim_ML_neuron_ExcType=1.txt','wt');

for I_stim=I_stim_min:dI_stim:I_stim_max

stat = @(V) (g_Ca*m_inf(V)*(-V + V_Ca) + g_K*w_inf(V)*(-V + V_K) + g_L*(-V + V_L) + I_stim);

V_start = -60; % mV 
format long
V_st = fzero(stat,V_start); % dV/dt = 0

k = k + 1;
I_stim_rec(k) = I_stim;
V_st_rec(k) = V_st;

a = w_inf(V_st);

b = (0.5./V4).*1/(cosh((V_st-V3)/V4))^2; % (mV)^(-1)

p = m_inf(V_st);

q = (0.5./V2).*1/(cosh((V_st-V1)/V2))^2; % (mV)^(-1)

A(k) = (g_Ca*(p + q*(V_st - V_Ca)) + g_K*a + g_L)/C_m; % kHz = 10^3 S/F

B(k) = g_K*b*(V_st - V_K)/C_m ;% kHz

tau(k) = tau_w(V_st)/beta; % ms

Y(k) = A(k) + 1./tau(k); % kHz

Z(k) = (A(k) + B(k))./tau(k); % (kHz)^2

gamma(k) = Y(k)/2; % kHz

omega_0(k) = sqrt(Z(k)); % kHz

if ((omega_0(k))^2 - (gamma(k))^2)>0 % omega = sqrt((omega_0)^2 - (gamma)^2)
    omega(k) = sqrt((omega_0(k))^2 - (gamma(k))^2); % kHz
    Im_omega(k) = 0;
else
    Im_omega(k) = sqrt(-(omega_0(k))^2 + (gamma(k))^2); % Im(omega) =  sqrt(-(omega_0)^2 + (gamma)^2)
    omega(k) = 0;
end;

fprintf(fid,'%10.12f %10.12f %10.12f %10.12f %10.12f %10.12f %10.12f %10.12f %10.12f %10.12f\n',I_stim_rec(k),A(k),I_stim_rec(k),B(k),I_stim_rec(k),1./tau(k),I_stim_rec(k),Y(k),I_stim_rec(k),Z(k));

fprintf(fid2,'%10.12f %10.12f %10.12f %10.12f %10.12f %10.12f %10.12f %10.12f\n',I_stim_rec(k),omega_0(k),I_stim_rec(k),gamma(k),I_stim_rec(k),omega(k),I_stim_rec(k),Im_omega(k));

if gamma(k)<0
    fprintf(fid3,'%10.12f %10.12f\n',I_stim_rec(k),gamma(k));
else
    fprintf(fid4,'%10.12f %10.12f\n',I_stim_rec(k),gamma(k));
end;

if Z(k)<0 % Z = (omega_0)^2
    fprintf(fid5,'%10.12f %10.12f\n',I_stim_rec(k),sqrt(-Z(k))); % Im(omega_0)
else
    fprintf(fid6,'%10.12f %10.12f\n',I_stim_rec(k),omega_0(k));
end;

if ((omega_0(k))^2 - (gamma(k))^2)<0 % omega = sqrt((omega_0)^2 - (gamma)^2)
    fprintf(fid7,'%10.12f %10.12f\n',I_stim_rec(k),Im_omega(k)); % Im(omega)
else
    fprintf(fid8,'%10.12f %10.12f\n',I_stim_rec(k),omega(k));
end;

(I_stim-I_stim_min)/(I_stim_max-I_stim_min)

end;

fclose(fid); fclose(fid2);       
        
figure(1)
plot(I_stim_rec,A,'blue',I_stim_rec,B,'red',I_stim_rec,1./tau,'green',I_stim_rec,Y,'black')
xlabel('I_{stim} [µA/cm^2]')
ylabel('A (blue), B (red), 1/\tau (green), Y (black) [kHz]')

figure(2)
plot(I_stim_rec,Z,'blue')
xlabel('I_{stim} [µA/cm^2]')
ylabel('Z [(kHz)^2]')

figure(3)
plot(I_stim_rec,(Z - (Y./2).^2),'blue')
xlabel('I_{stim} [µA/cm^2]')
ylabel('omega^{2} [(kHz)^2]')

figure(4)
plot(I_stim_rec,omega_0,'blue',I_stim_rec,gamma,'red',I_stim_rec,omega,'black')
xlabel('I_{stim} [µA/cm^2]')
ylabel('omega_{0} (blue), gamma (red), omega (black) [kHz]')

figure(5)
plot(I_stim_rec,V_st_rec,'blue')
xlabel('I_{stim} [µA/cm^2]')
ylabel('V_{st} [mV]')
```
