## Supplementary material for "Analytical solution of linearized equations of the Morris-Lecar neuron model at large constant stimulation": Two supplementary Figures, MATLAB scripts, and generated data used for Figures: ReadMe.pdf

The difference between "Less accurate but faster code" (ML_neuron_damped_lin_eq_parameters_of_I_stim.m) and "More accurate but slower code" (ML_neuron_damped_lin_eq_parameters_of_I_stim_accurate.m) is only that in the last case the numerical instability in the vicinity of I_min = 40 μA/cm$^2$ has been eliminated by additionally using **fsolve**, which can find all equation roots, rather than the nearest root to the starting value, as **fzero** does.
