## Supplementary material for "Analytical solution of linearized equations of the Morris-Lecar neuron model at large constant stimulation": Two supplementary Figures, MATLAB scripts, and generated data used for Figures: ML_neuron_freq_ode23s_tau_w_equal_tau_of_V_st.m

```
clear all;

% potential V is in mV
% time t is in ms
% electical capacity C_m is in µF/cm^2
% conductivity g is in mS/cm^2
% stimulating current I_stim is in µA/cm^2

% Parameters for the Morris-Lecar model were taken from J. Rinzel, B. Ermentrout, Analysis of neural excitability and oscillations. 
% In: C. Koch, I. Segev (Eds.), Methods in neuronal modeling: From synapses to networks (2nd Ed., MIT Press, Cambridge, 1998), pp. 251-291.

C_m = 20; % µF/cm^2

g_K = 8; % mS/cm^2
g_L = 2; % mS/cm^2

V_Ca = 120; % mV
V_K = -84; % mV
V_L = -60; % mV
V1 = -1.2; % mV
V2 = 18; % mV

excitability_type = 1;

if (excitability_type == 1)
    g_Ca = 4; % mS/cm^2
    V3 = 12; % mV
    V4 = 17.4; % mV
    beta = 0.067; % kHz, beta = 1/tau_max, tau_max = 14.925 ms
    tau_max = 1/beta; % ms
end;

if (excitability_type == 2)
    g_Ca = 4.4; % mS/cm^2
    V3 = 2; % mV
    V4 = 30; % mV
    beta = 0.04; % kHz, beta = 1/tau_max, tau_max = 25 ms
    tau_max = 1/beta; % ms
end;

%-------------- Functions------------------

m_inf = @(V) 0.5.*(1 + tanh((V-V1)./V2)); 
w_inf = @(V) 0.5.*(1 + tanh((V-V3)./V4)); 
tau_w = @(V) 1./cosh((V-V3)./(2.*V4)); 

rest = @(V) (g_Ca*m_inf(V)*(-V + V_Ca) + g_K*w_inf(V)*(-V + V_K) + g_L*(-V + V_L));

%---------- Finding V_rest  ----------- %
V_start = -60; % mV 
format long
V_rest = fzero(rest,V_start) % sum of ion currents = 0
rest(V_rest)
%--------------------------%

V_th = 20.0; % mV, spike-detection threshold 

% I_stim is constant stimulating current, µA/cm^2
I_stim_min = 110;
dI_stim = 0.1;
I_stim_max = 150;

t_max = 20000; % ms;
time = [0 t_max];
k = 0;
freq = zeros(1,1);
I_stim_rec = zeros(1,1);
tau_rec = zeros(1,1);

fid = fopen('freq_of_I_stim_ML_neuron_ode23s_ExcType=1_tau_w=tau(V_st).txt','wt');
fid2 = fopen('tau(V_st)_of_I_stim_ML_neuron_ode23s_ExcType=1.txt','wt');

for I_stim=I_stim_min:dI_stim:I_stim_max

spikes = 0; 
mark = 0; 

%---------- Finding V_st and tau(V_st) ----------- %
V_start = 10; % mV 
format long
V_st = fzero(@(V)(g_Ca*m_inf(V)*(-V + V_Ca) + g_K*w_inf(V)*(-V + V_K) + g_L*(-V + V_L) + I_stim),V_start);
tau = tau_w(V_st)/beta; % ms
%--------------------------%

% Below V = y(1) and w = y(2):

F = @(t,y) [(g_Ca.*m_inf(y(1)).*(-y(1) + V_Ca) + g_K.*y(2).*(-y(1) + V_K) + g_L.*(-y(1) + V_L) + I_stim)./C_m; (w_inf(y(1))-y(2))./tau]; 

[T, Y] = ode23s(F,time,[V_rest w_inf(V_rest)]);

V = Y(:,1);

for i=1:length(V)

 if (V(i)>V_th) 
   mark = 1;
 end;                            
 
 if (mark>0)&&(V(i)<0)
    mark=-1;
	spikes = spikes + 1;
 end;
  
end;

k = k + 1;
I_stim_rec(k) = I_stim;
tau_rec(k) = tau;
if spikes>1
freq(k) = spikes/(t_max/1000);
else
freq(k) = 0;    
end;
fprintf(fid,'%10.12f %10.12f\n',I_stim_rec(k),freq(k));
fprintf(fid2,'%10.12f %10.12f\n',I_stim_rec(k),tau_rec(k));
I_stim/I_stim_max

end;

fclose(fid); 
fclose(fid2); 
        
figure(1)
plot(I_stim_rec,freq,'b','Marker','.')
xlabel('I_{stim} [µA/cm^2]')
ylabel('Average spiking frequency at tau(V)=tau(V_{st}) [Hz]')

figure(2)
plot(I_stim_rec,tau_rec./tau_max,'b','Marker','.')
xlabel('I_{stim} [µA/cm^2]')
ylabel('tau(V_{st})/tau_{max}')
```
