## Supplementary material for "Analytical solution of linearized equations of the Morris-Lecar neuron model at large constant stimulation": Two supplementary Figures, MATLAB scripts, and generated data used for Figures: Figure_S1.pdf

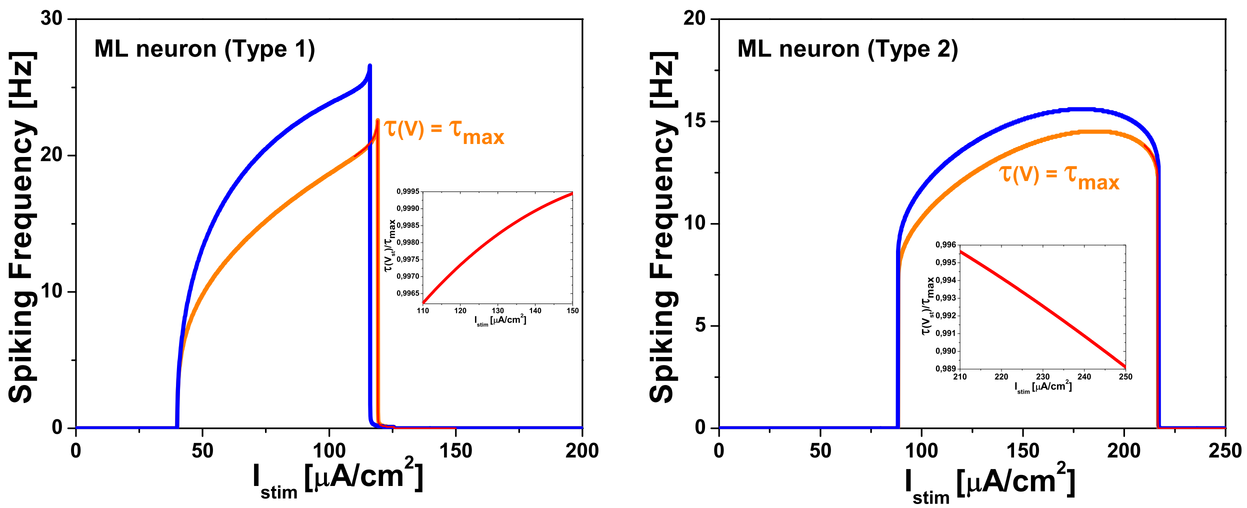


**Figure S1**. Dependence of spike generation frequency (determined as the number of spikes divided by the time interval of 20000 ms) on constant stimulating current $I_{stim}$ for the Morris-Lecar (ML) model with the 1st (left graph) and 2nd (right graph) excitability types. The blue curves correspond to the standard ML model described in Sec. 2 of the main text. The orange curves correspond to a simplified version of the ML model with function $\tau(V)$, see Eq. (5), taken as a constant equal to its maximal value $\tau_{\max}$. In turn, the red curves near the upper boundary of the sustained spiking interval, which are virtually superimposed on the orange ones, correspond to the similar case where function $\tau(V)$ is also taken as a constant equal to $\tau(V_{st})$. The value $V_{st}$ is determined from equation $I_{ion}(V_{st}, w_\infty(V_{st})) = I_{stim}$ and, above the upper boundary, $V_{st}$ corresponds to the stationary asymptotic value of the neuron potential. Finally, the inset in each graph shows the dependence of $\tau(V_{st})$ on $I_{stim}$ that is practically negligible (nevertheless, note that it is opposite for type 1 and type 2) so that one can safely use universal approximation $\tau(V) = \tau_{\max}$.

The parameters of the ML model with the excitability type 1 were as follows: $C_m = 20$ $\mu$F/cm$^2$, $g_{Ca} = 4$ mS/cm$^2$, $g_K = 8$ mS/cm$^2$, $g_L = 2$ mS/cm$^2$, $V_{Ca} = 120$ mV, $V_K = -84$ mV, $V_L = -60$ mV, $V_1 = -1.2$ mV, $V_2 = 18$ mV, $V_3 = 12$ mV, $V_4 = 17.4$ mV, $\tau_{\max} = 14.925$ ms. These parameters result in the following values for the lower and upper boundaries of the sustained spiking interval of $I_{stim}$: $I_{min} = 40$ $\mu$A/cm$^2$ and $I_{max} = 116.1$ $\mu$A/cm$^2$.

In turn, the ML model of the excitability type 2 had the following parameters: $g_{Ca} = 4.4$ mS/cm$^2$, $V_3 = 2$ mV, $V_4 = 30$ mV, $\tau_{\max} = 25$ ms, with all the rest parameters being the same as those for the type 1. The corresponding values for the lower and upper boundaries of the sustained spiking interval are $I_{min} = 88.3$ $\mu$A/cm$^2$ and $I_{max} = 216.9$ $\mu$A/cm$^2$.
