## Supplementary material for "Analytical solution of linearized equations of the Morris-Lecar neuron model at large constant stimulation": Two supplementary Figures, MATLAB scripts, and generated data used for Figures: Figure_S2.pdf

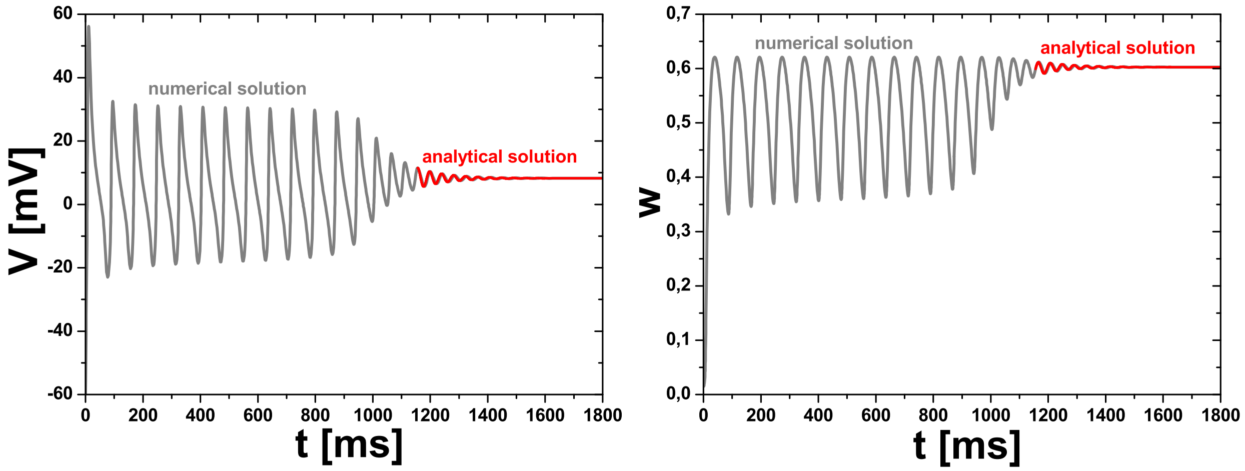


**Figure S2**. Left graph: The gray curve is a numerical solution for dynamics of neuron potential $V(t)$ in the Morris-Lecar (ML) model with the 2nd excitability type at $I_{stim} = 216.995\ \mu\text{A/cm}^2 > I_{max} = 216.9\ \mu\text{A/cm}^2$, where $I_{max}$ is the upper boundary of the sustained spiking interval (see the right graph in Fig. S1). The red curve is the analytical solution of the linearized system of equations of the ML model with initial conditions taken at the point of a local maximum of the potential ($t_0 = 1156$ ms, $V_0 = 11.49$ mV). Right graph: The corresponding numerical (gray) and analytical (red) solutions for $w(t)$. Parameters of the analytical formulas for this example are as follows: $V_{st} = 8.25$ mV, $a = 0.6$, $\omega_0 = 151.2$ Hz, $\gamma = 9.76$ Hz, $\omega = 150.9$ Hz, $1/\tau = 40.2$ Hz, $\eta = 0.065$, $\chi = 0.2$, $2\gamma/|A_K| = 14.43$, $U_0 = 3.24$ mV, $a/w_0 = 1.0$, $W_a = 0.014$, and $W_c = -0.005$, where the last five parameters depend on $t_0$ and $V_0$ values.

The ML model parameters for the 2nd neuronal excitability type were as follows: $C_m = 20$ $\mu\text{F/cm}^2$, $g_{Ca} = 4.4$ mS/cm$^2$, $g_K = 8$ mS/cm$^2$, $g_L = 2$ mS/cm$^2$, $V_{Ca} = 120$ mV, $V_K = -84$ mV, $V_L = -60$ mV, $V_1 = -1.2$ mV, $V_2 = 18$ mV, $V_3 = 2$ mV, $V_4 = 30$ mV, $\tau_{\max} = 25$ ms.
