## Supplementary material for "Analytical solution of linearized equations of the Morris-Lecar neuron model at large constant stimulation": Two supplementary Figures, MATLAB scripts, and generated data used for Figures: ReadMe.pdf

The MATLAB script ML_neuron_dynamics_ode23s.m, using built-in ode23s solver for stiff systems of ordinary differential equations, solves dynamic equations for V(t) and w(t) of the standard Morris-Lecar model for the 1st or 2nd excitability type (flag excitability_type = 1 or 2, respectively).

In turn, the MATLAB script ML_neuron_freq_ode23s.m finds the dependence of sustained spike generation frequency on constant stimulating current I_stim for the standard Morris-Lecar model of chosen excitability type.
