## Supplementary figures and images for "Analytical solution of linearized equations of the Morris-Lecar neuron model at large constant stimulation"

### figure(1).png

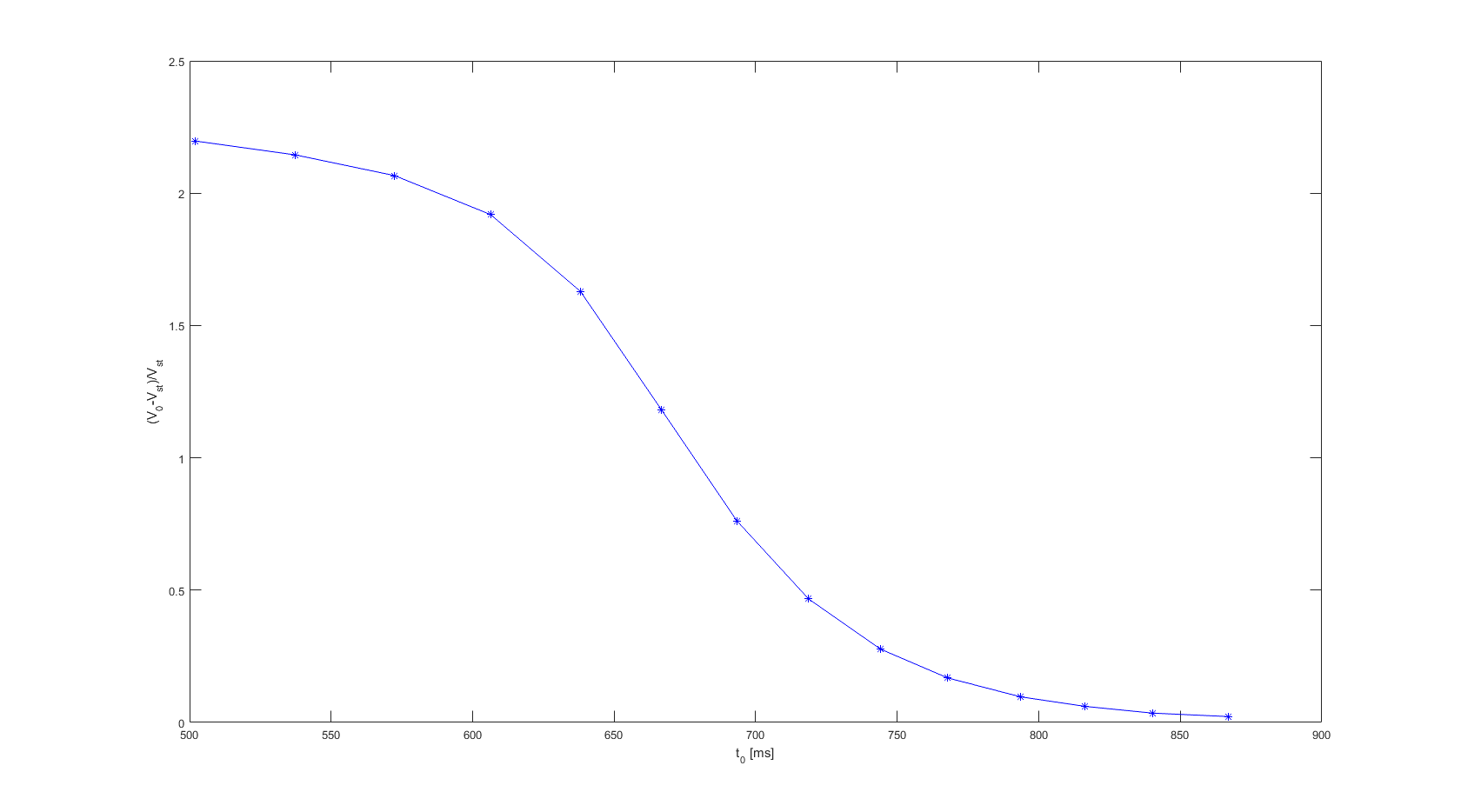

### figure(1).png

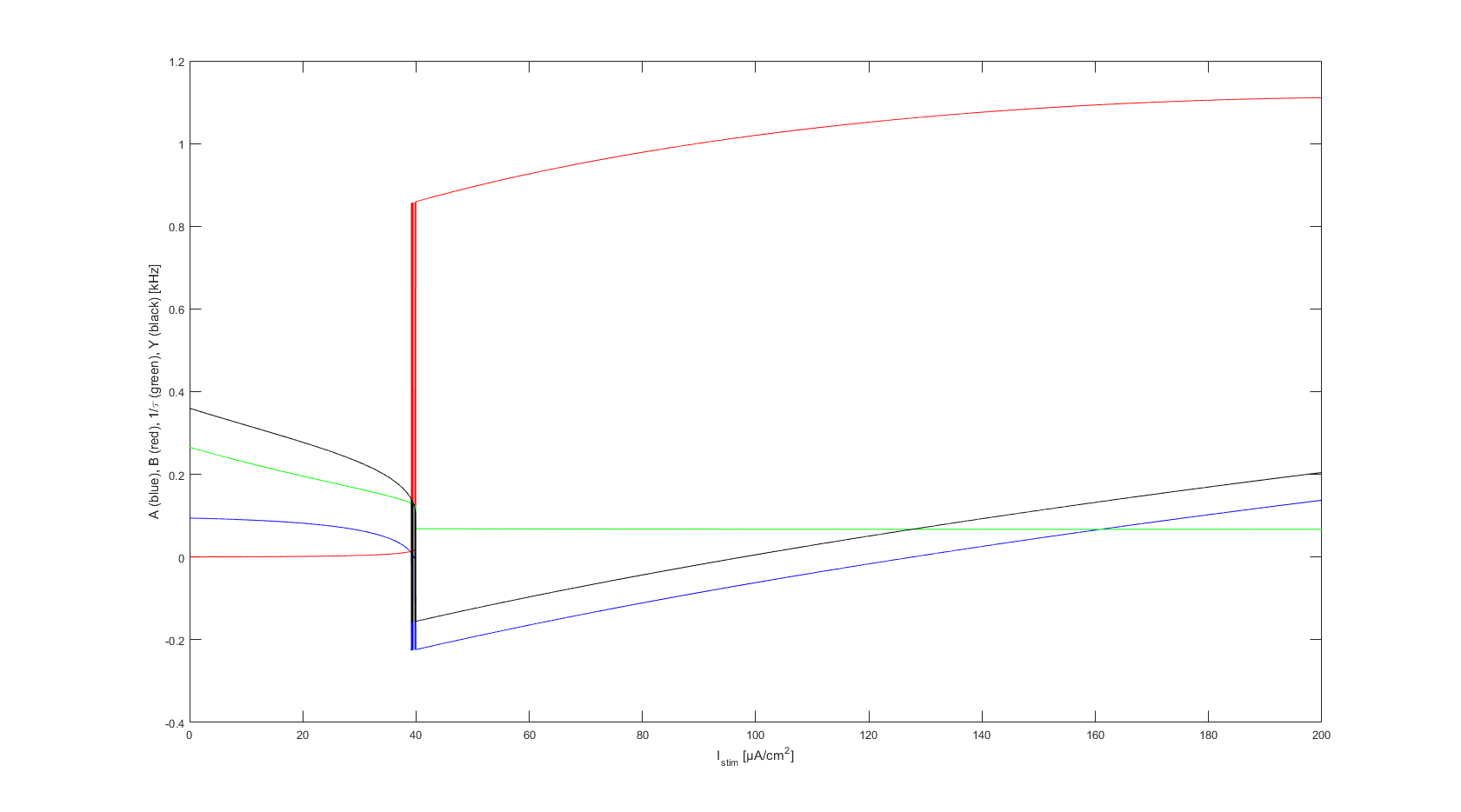

### figure(1).png

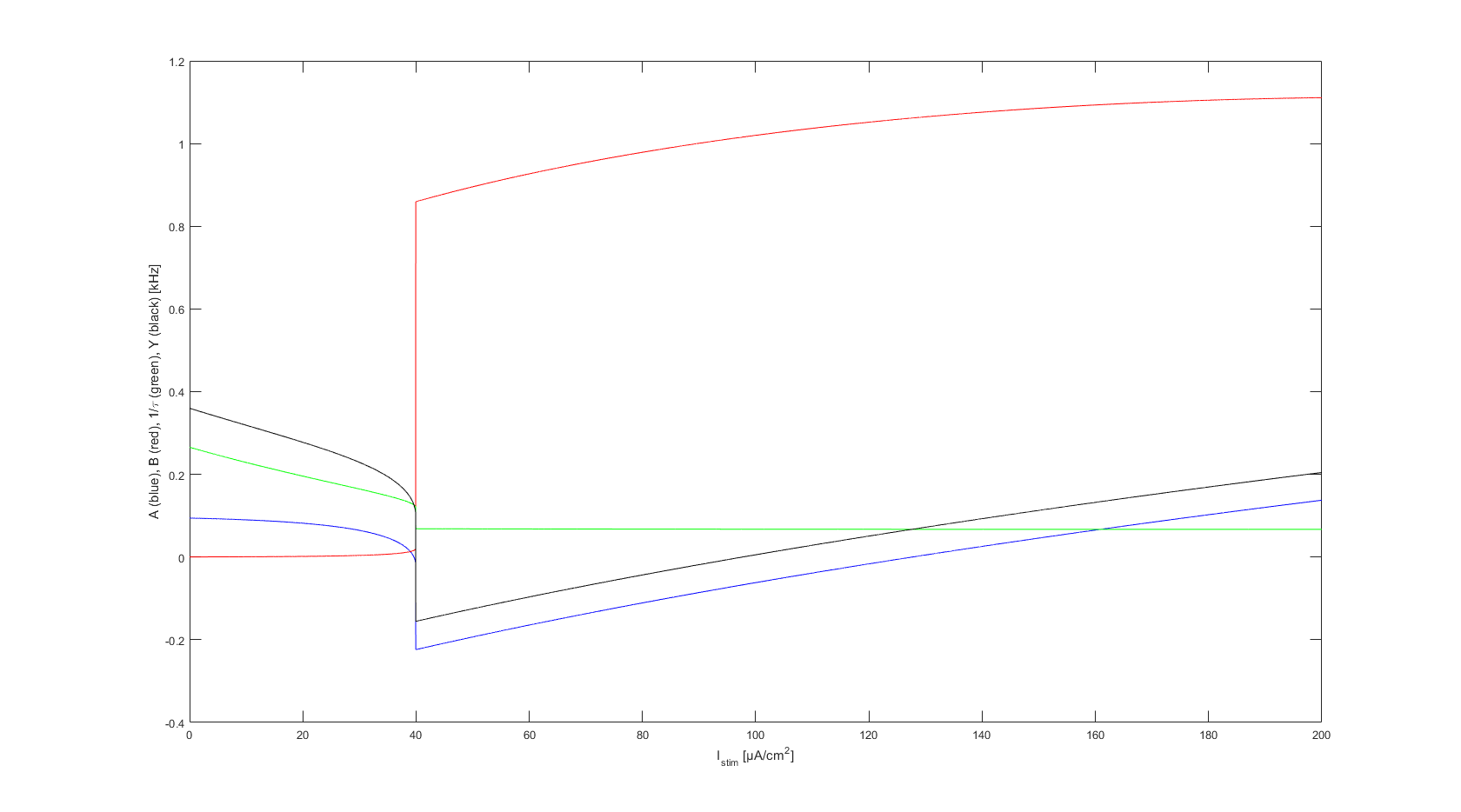

### figure(2).png

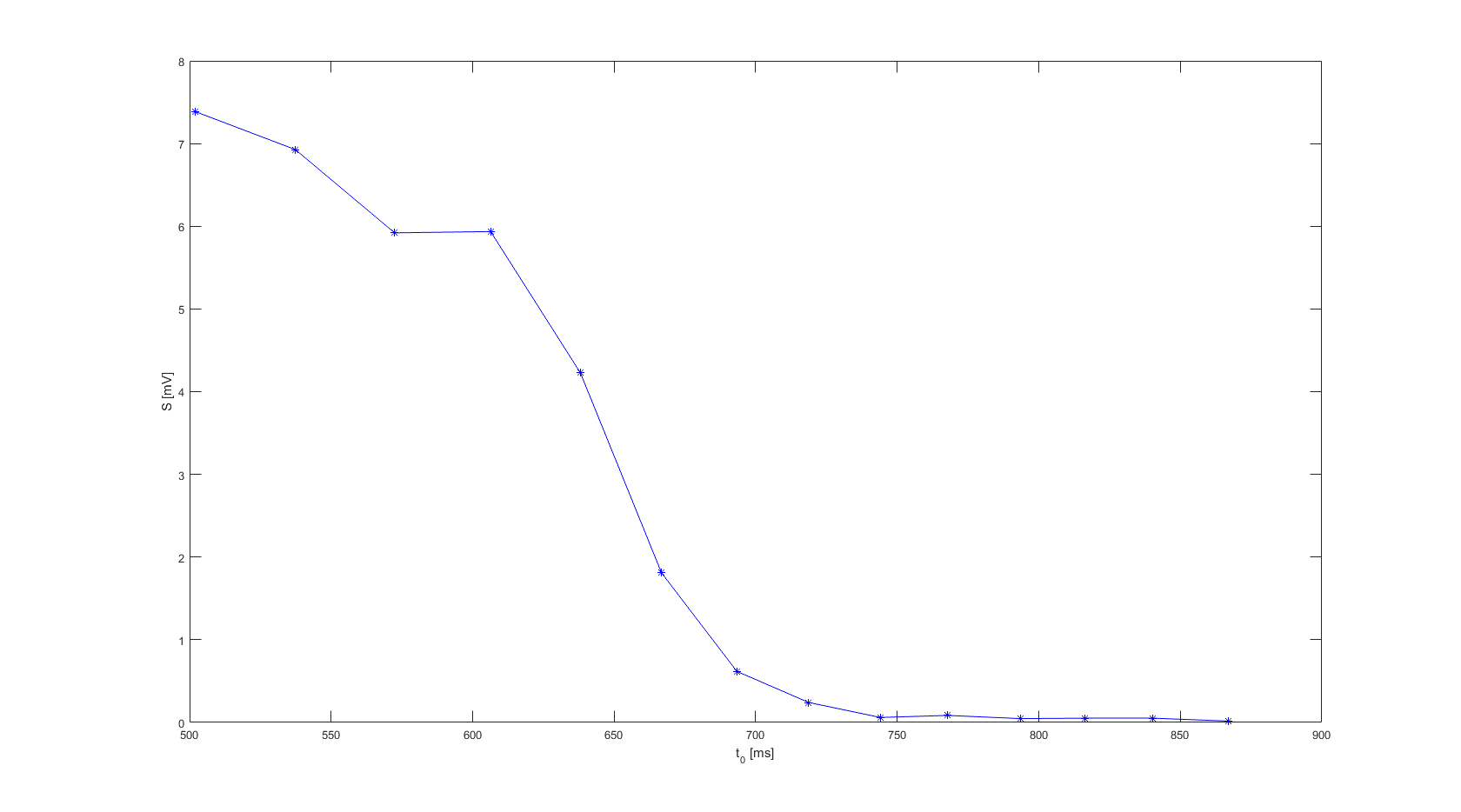

### figure(2).png

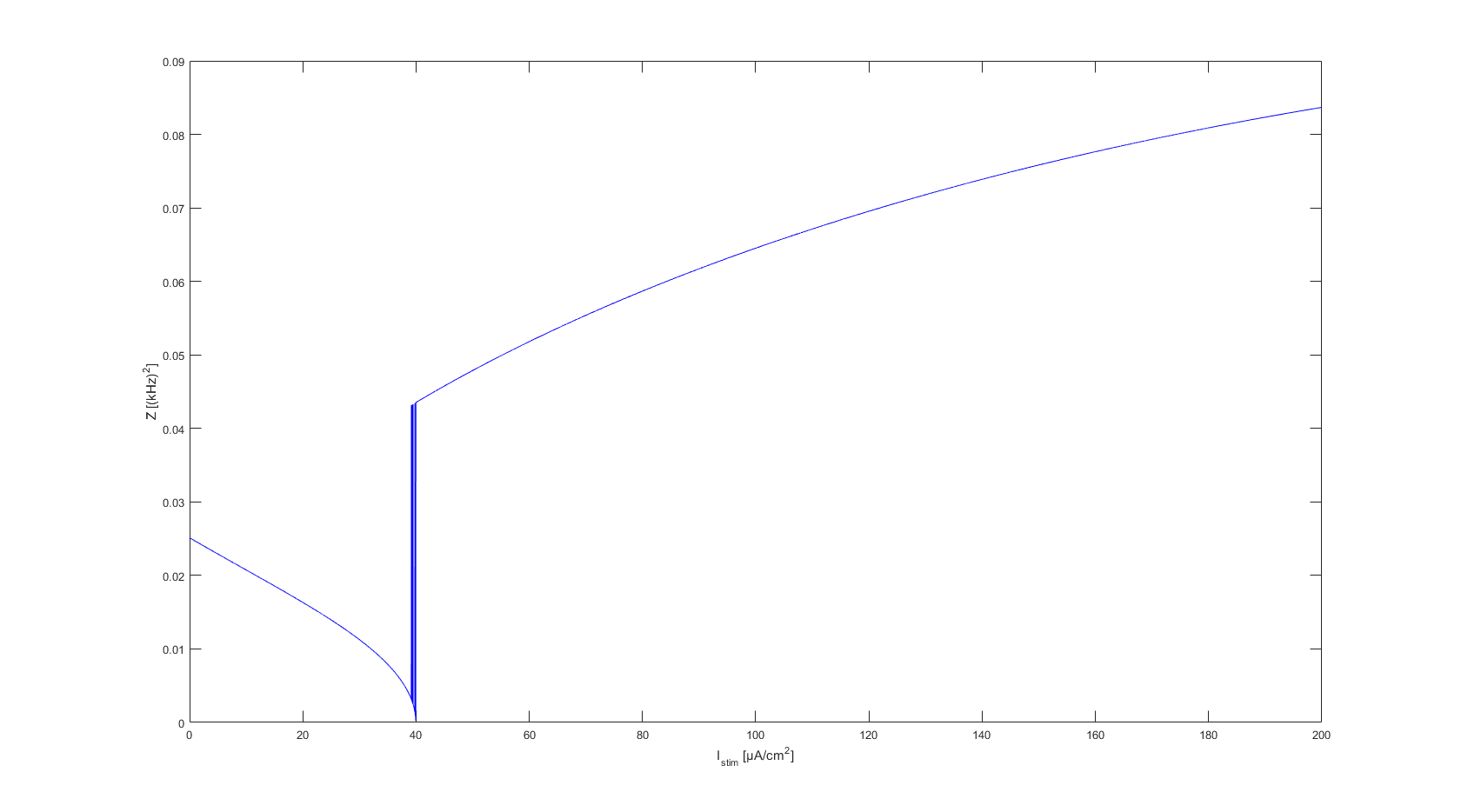

### figure(2).png

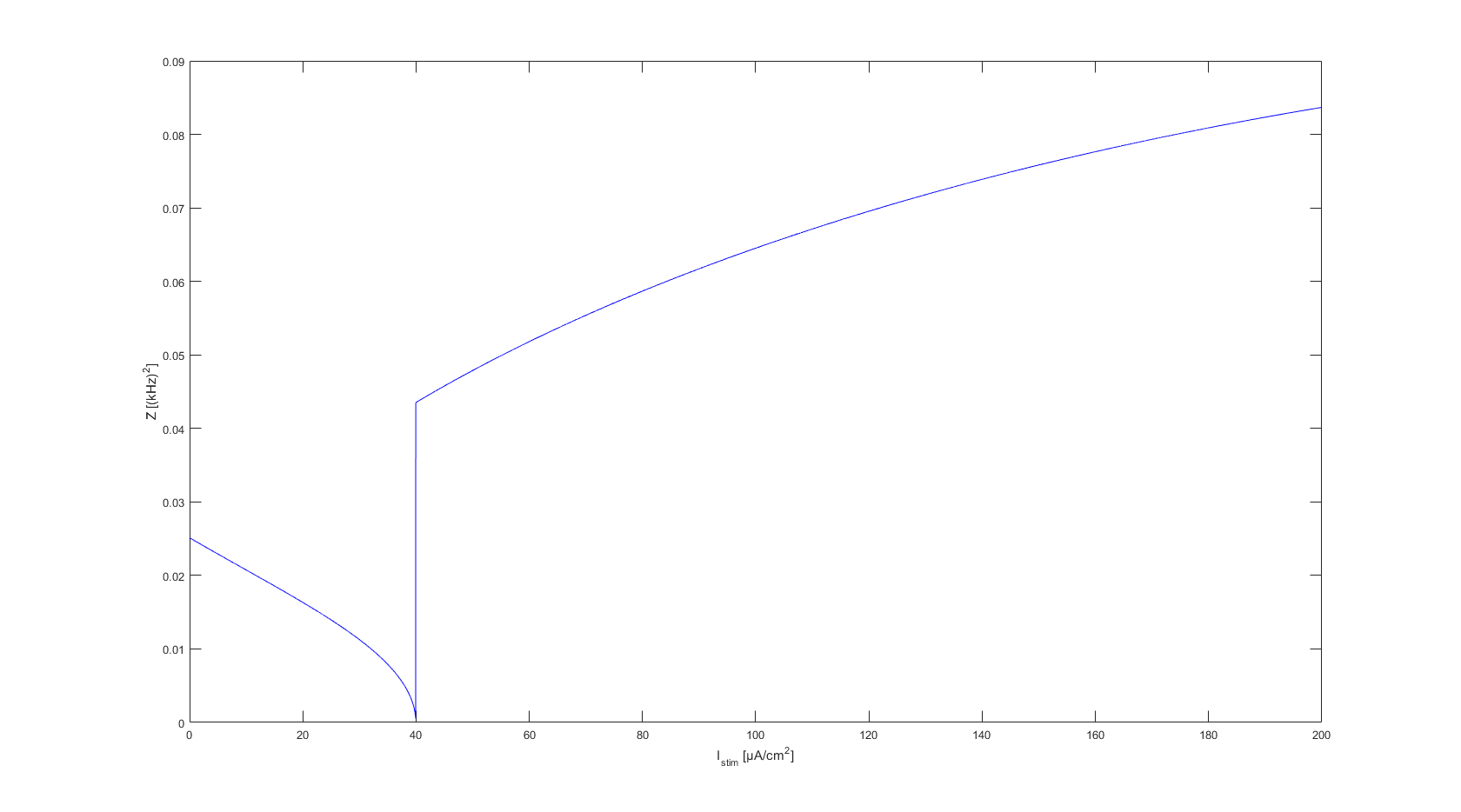

### figure(3).png

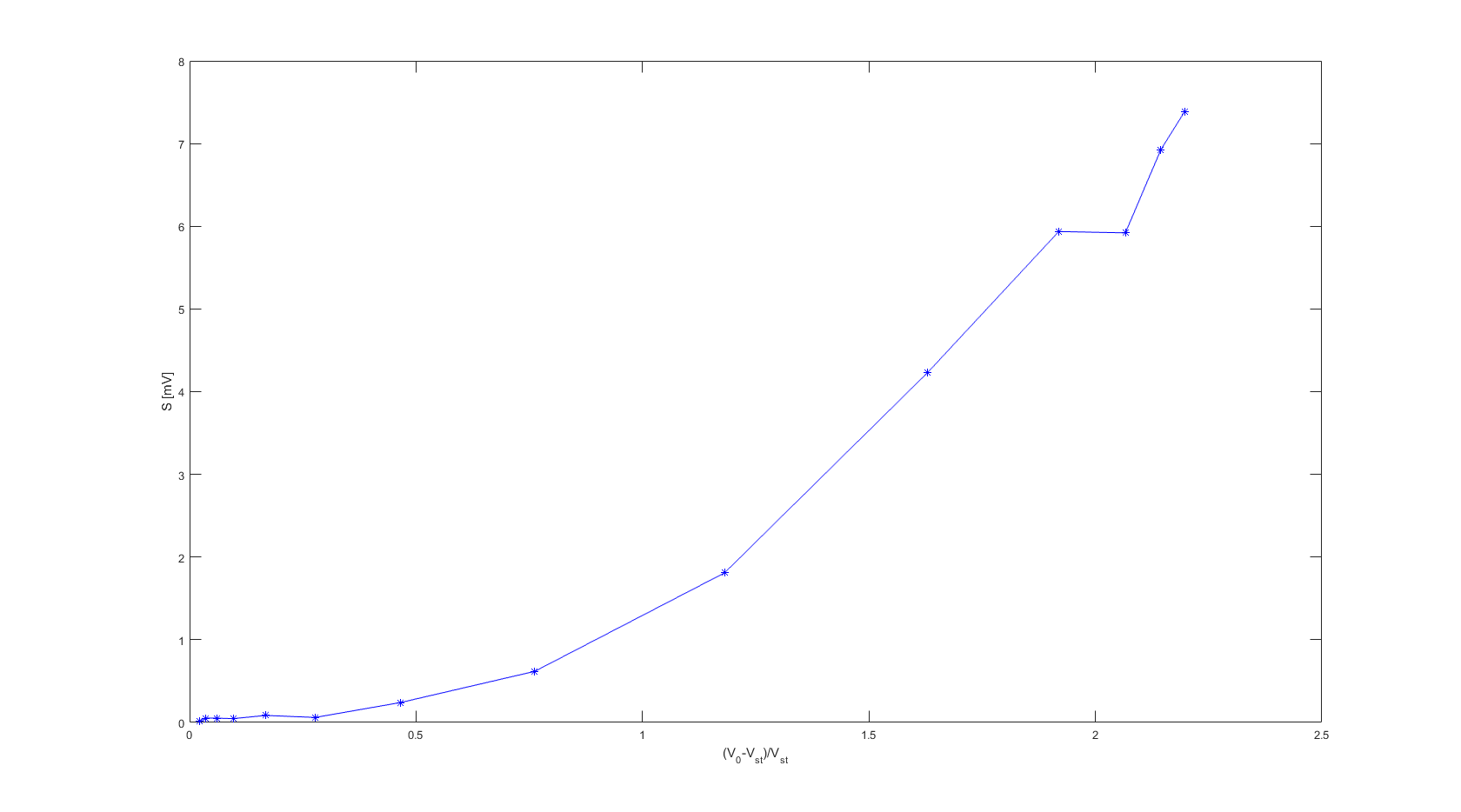

### figure(3).png

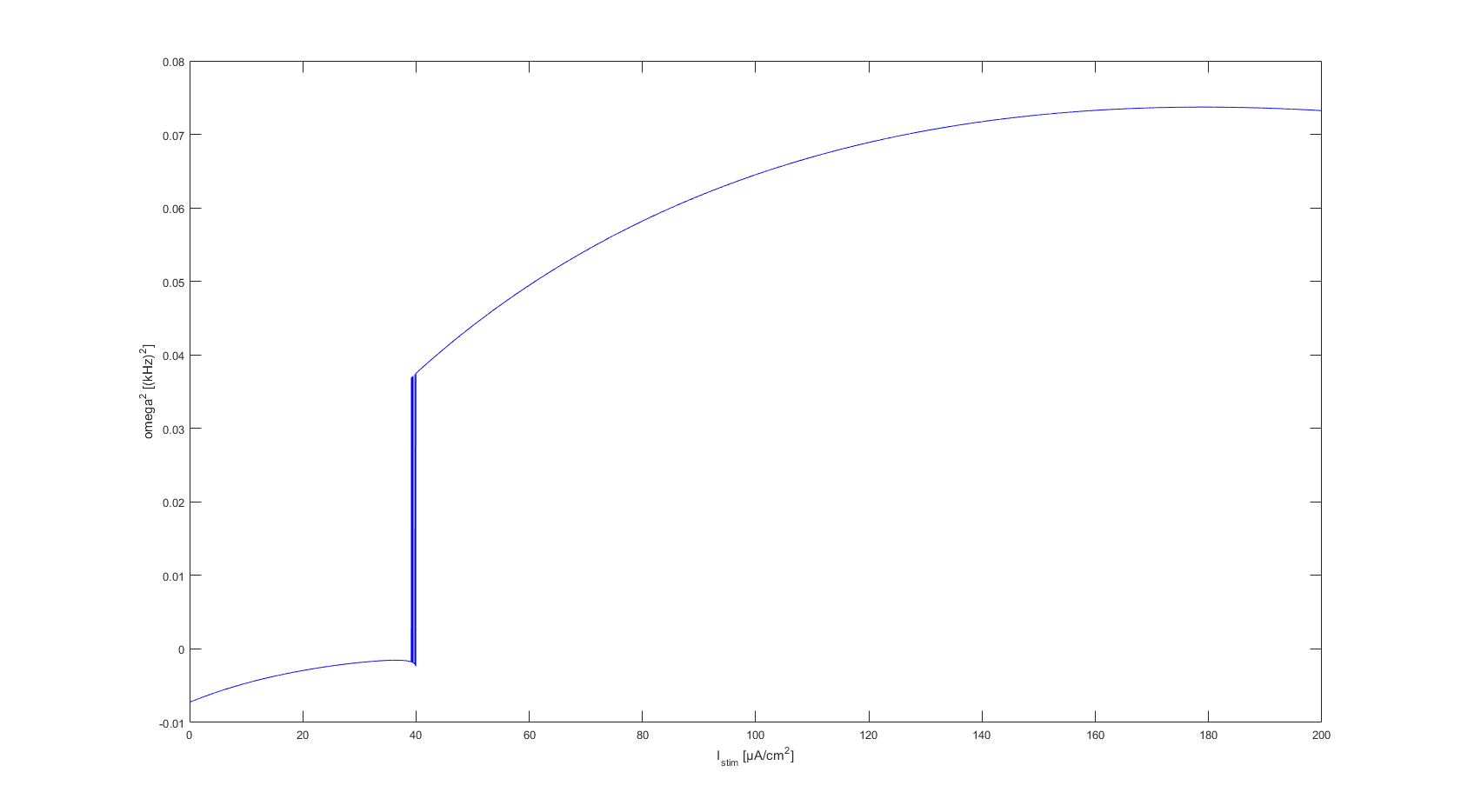

### figure(3).png

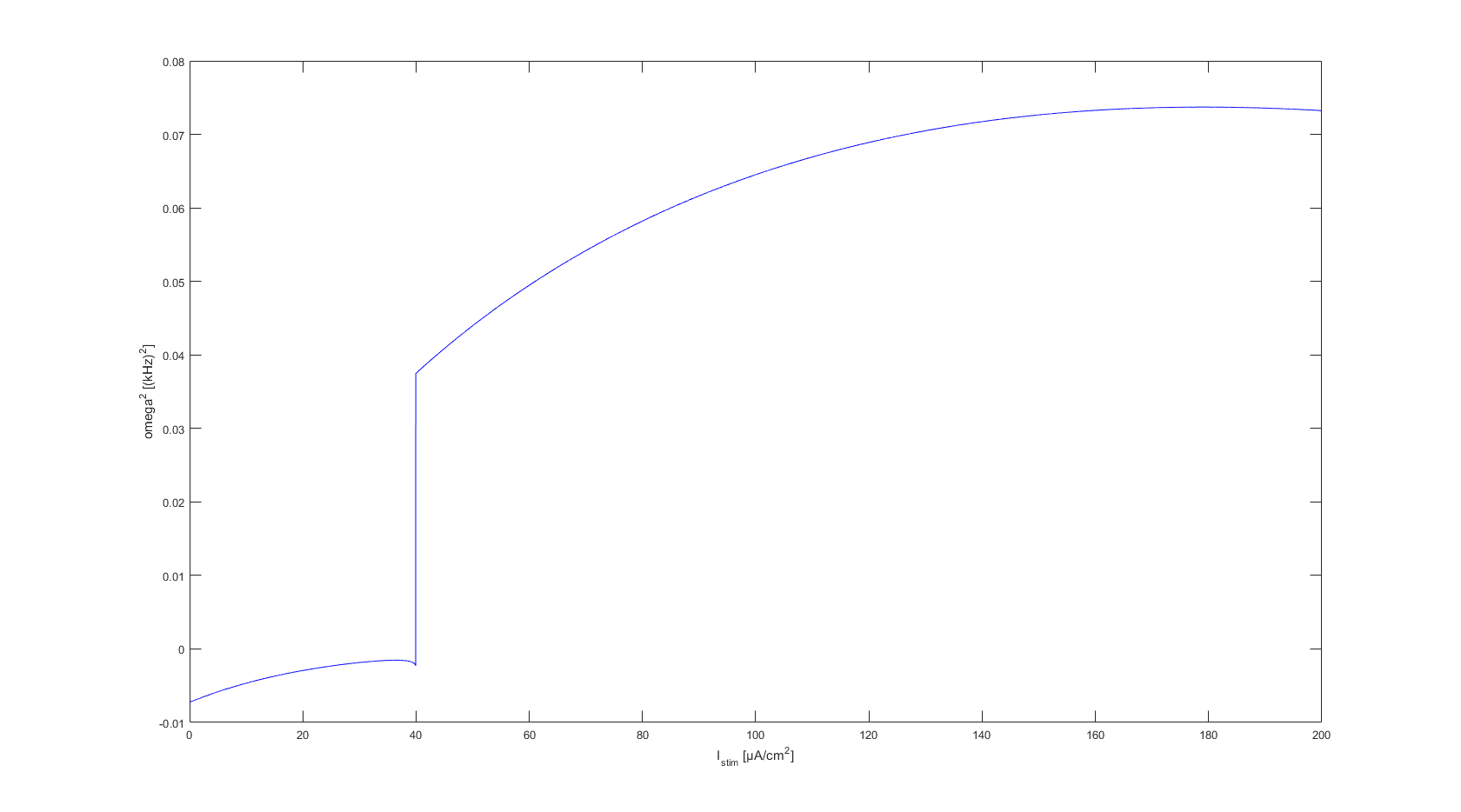

### figure(4).png

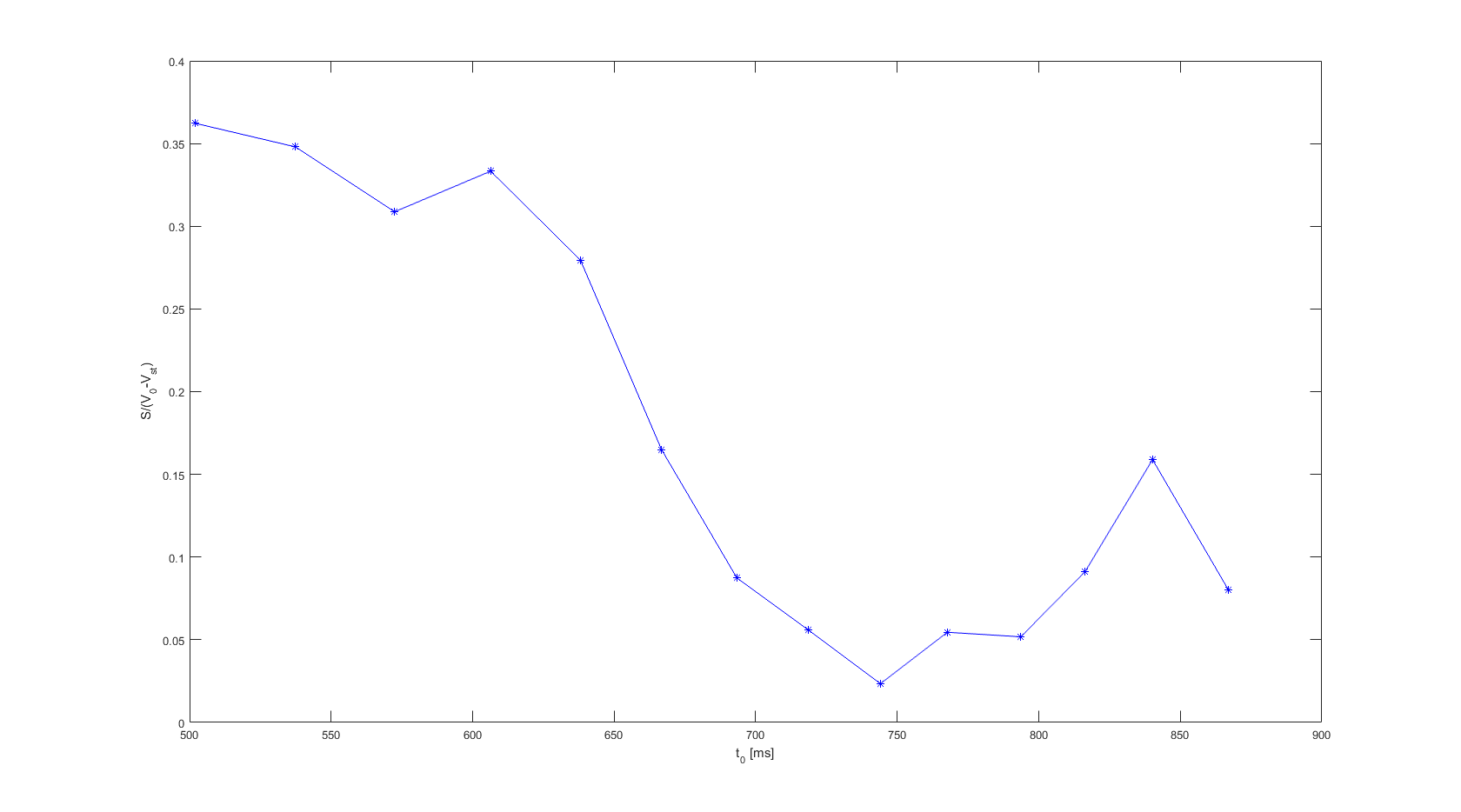

### figure(4).png

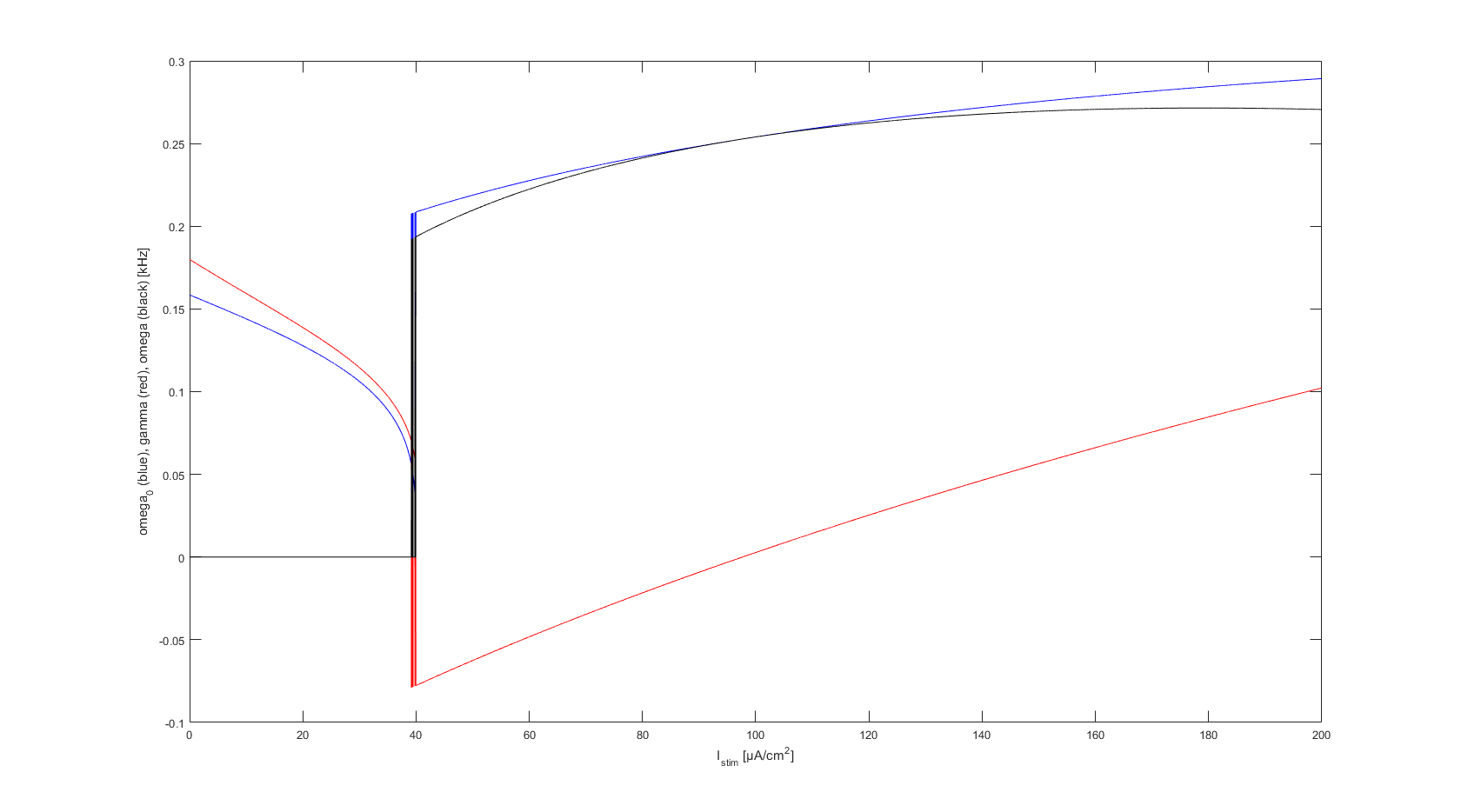

### figure(4).png

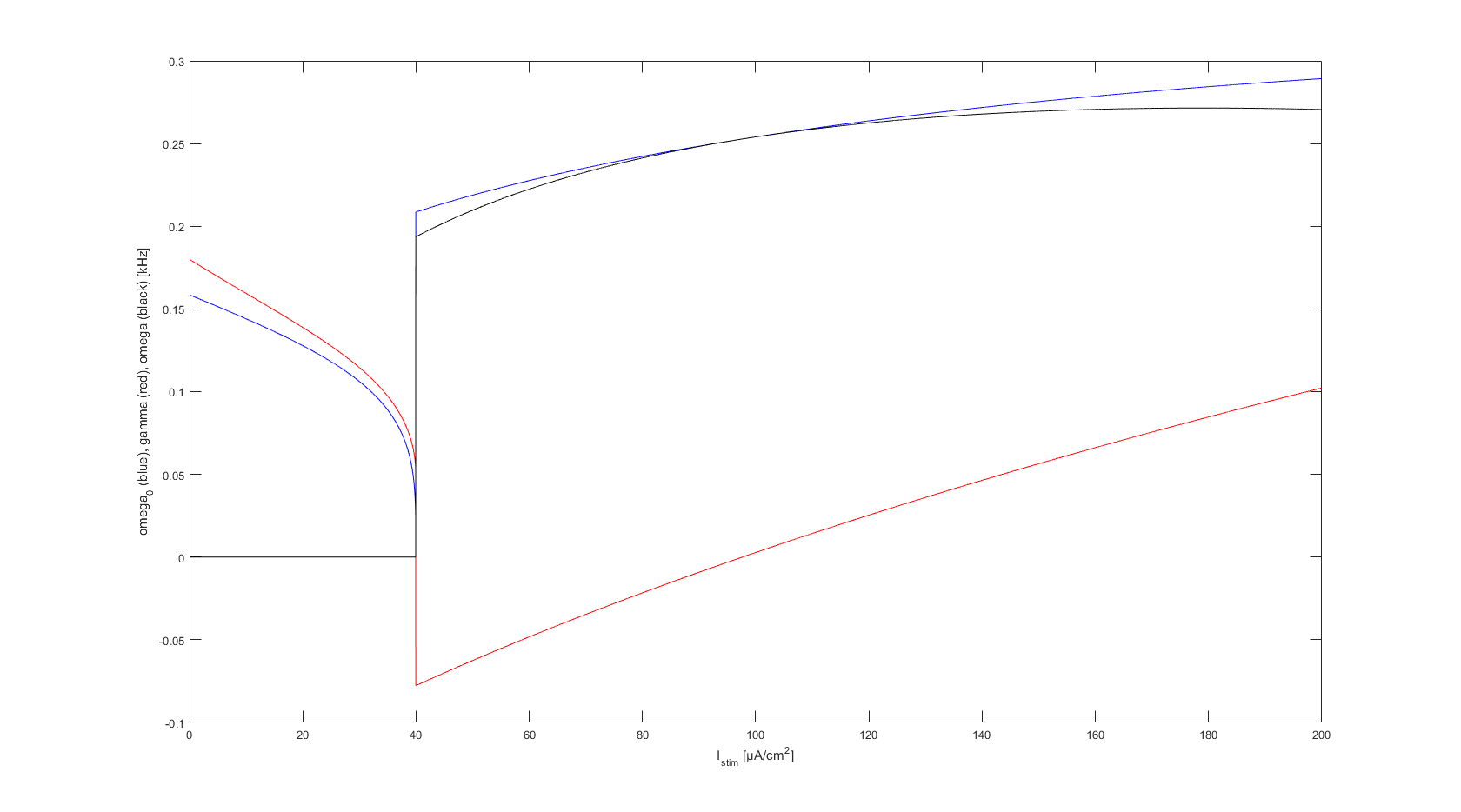

### figure(5).png

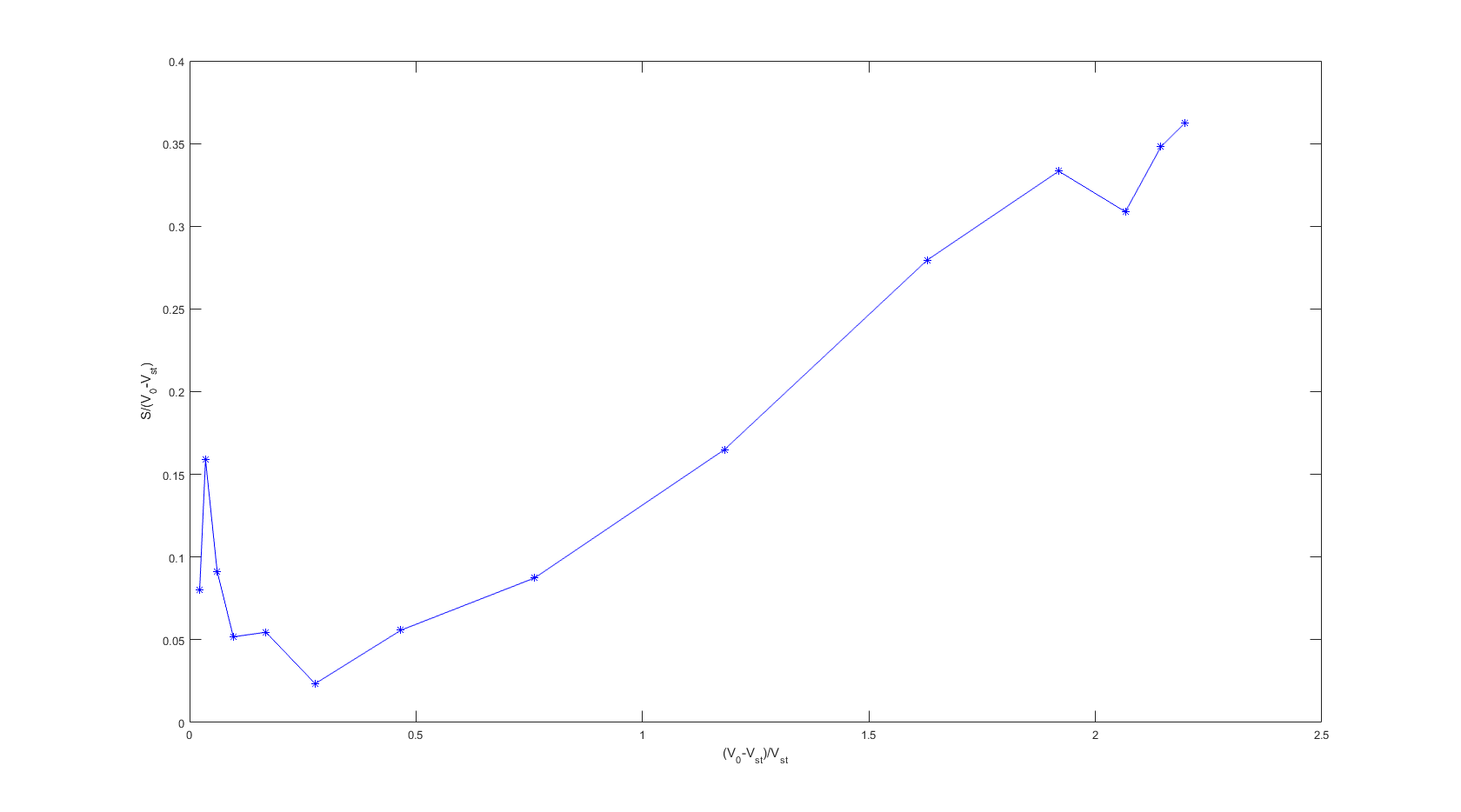

### figure(5).png

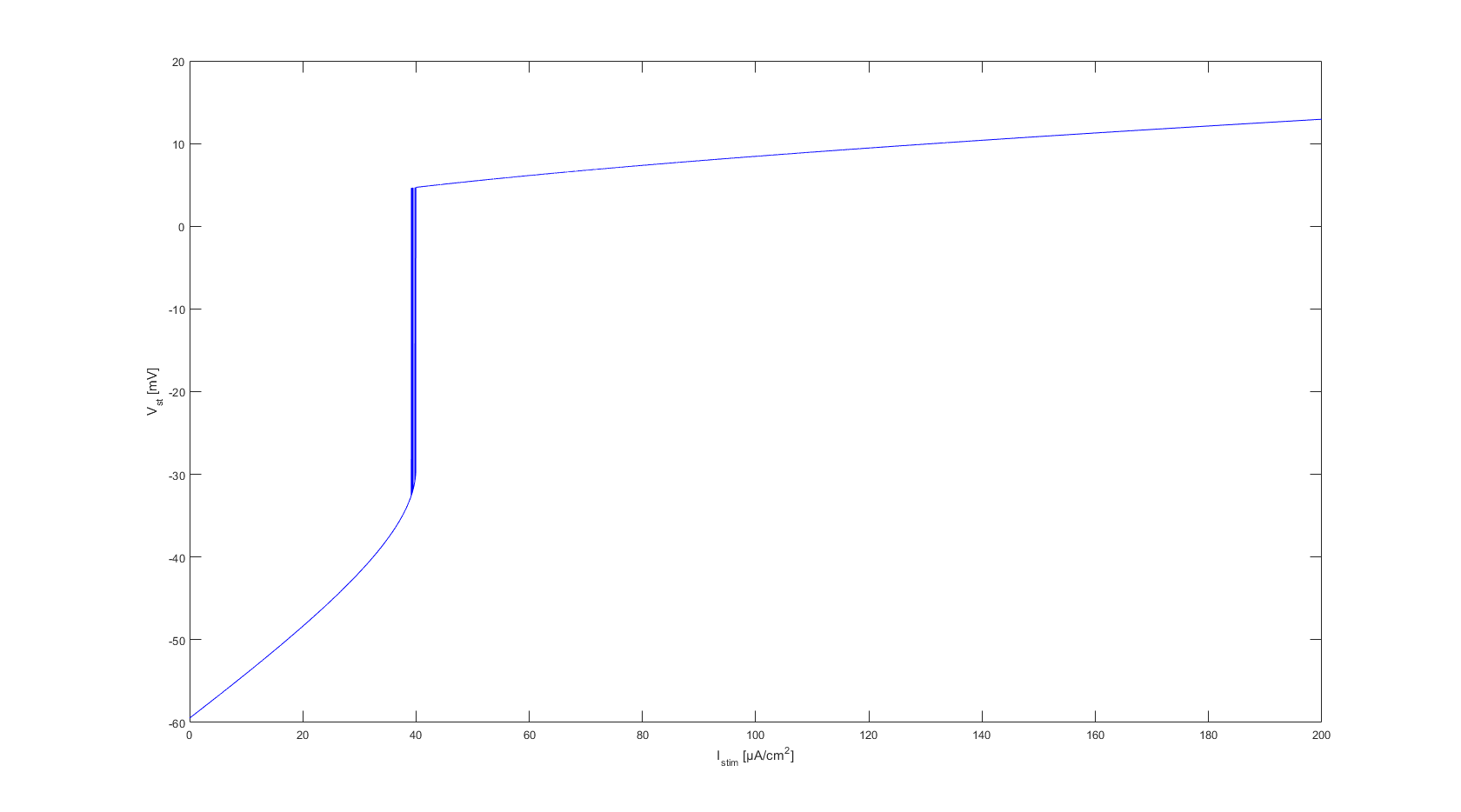

### figure(5).png

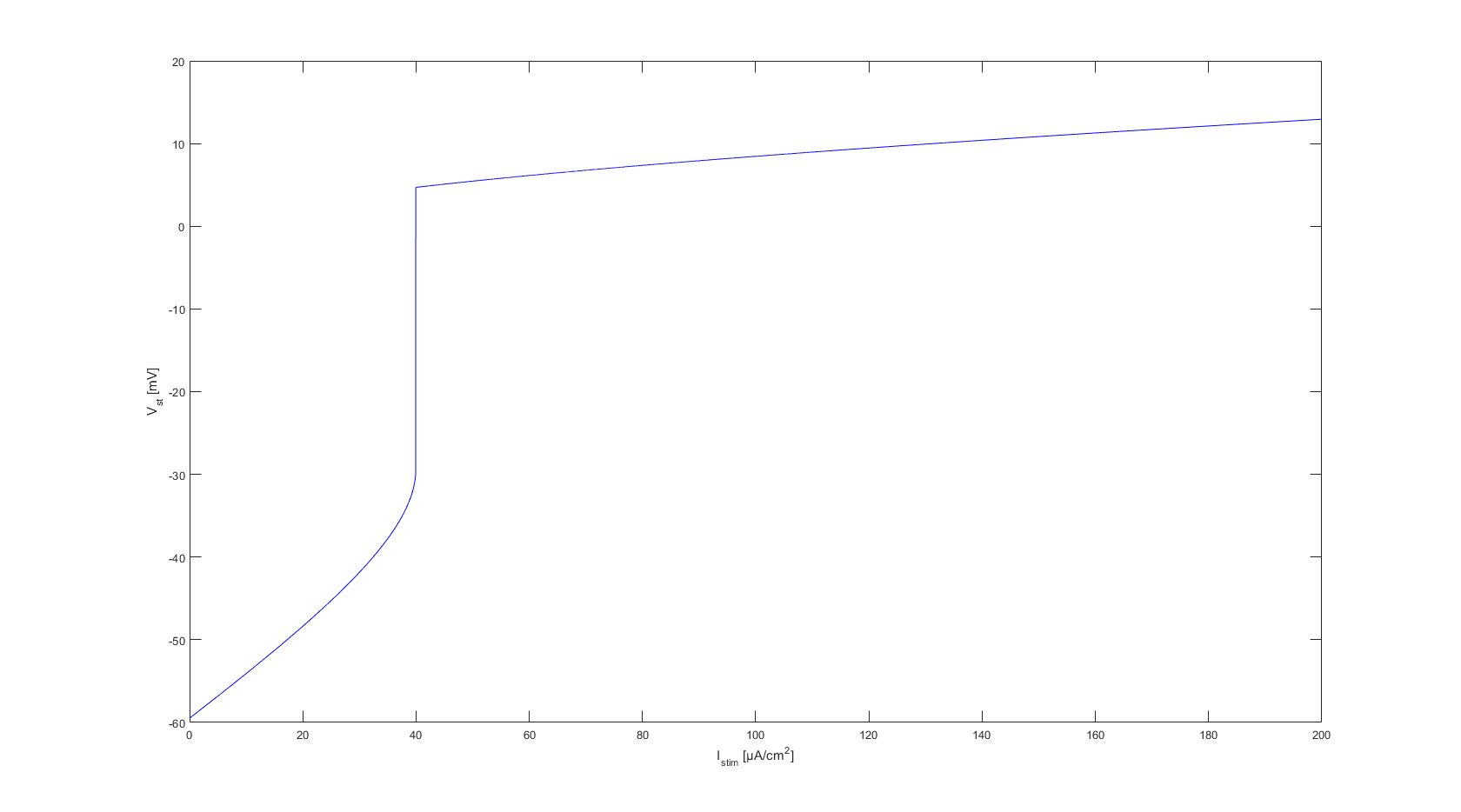

### figure(6).png

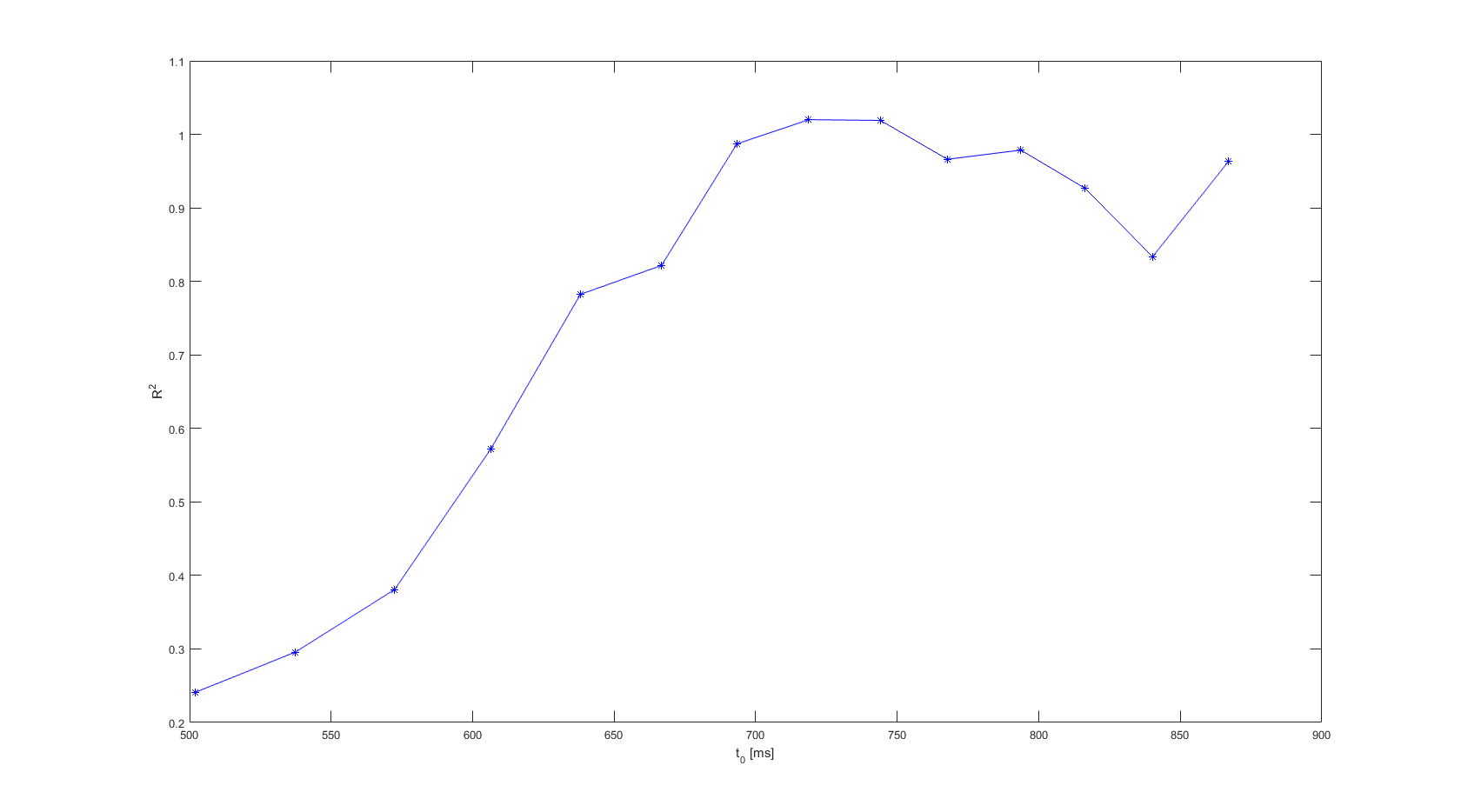

### figure(7).png

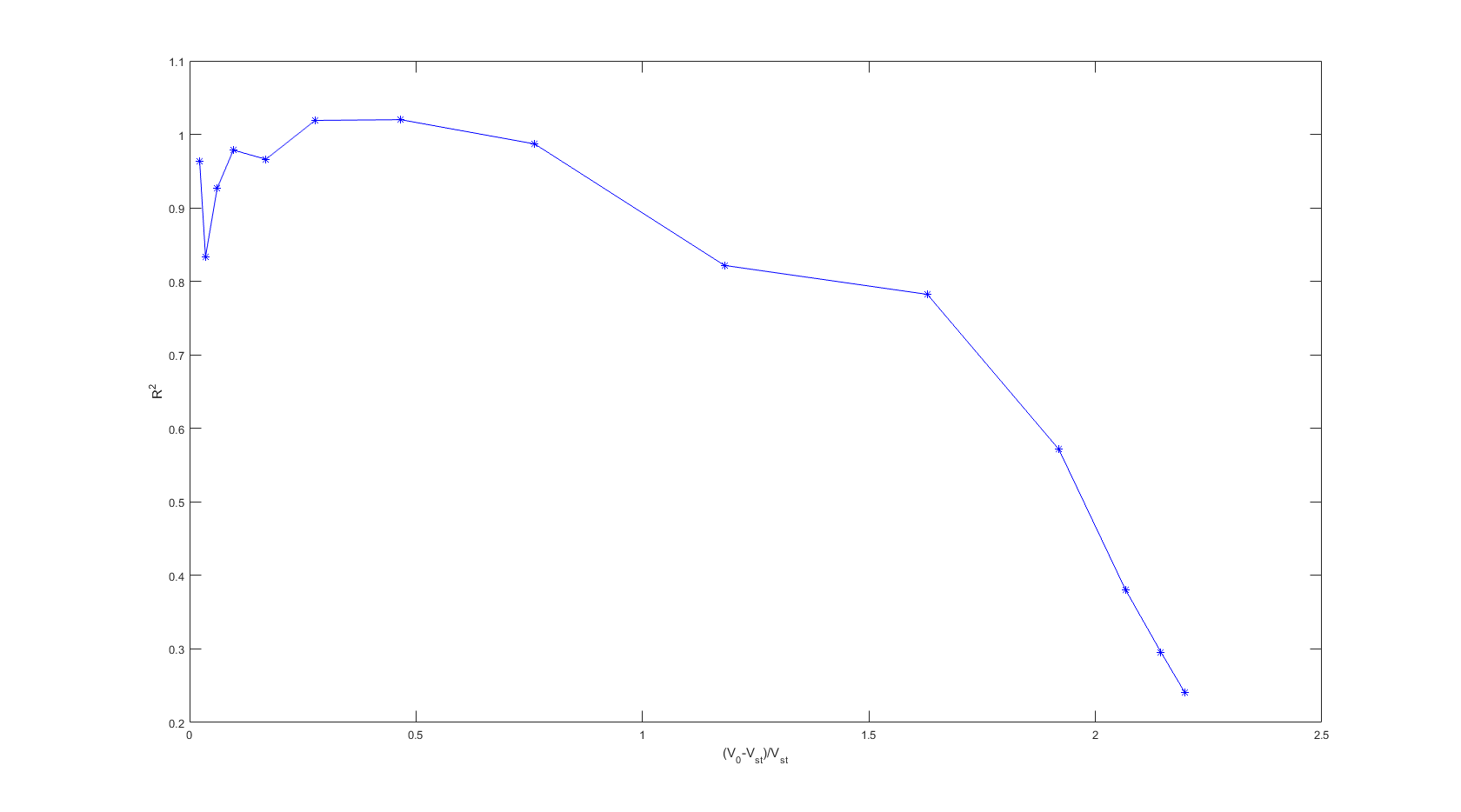

### Figure_1.png

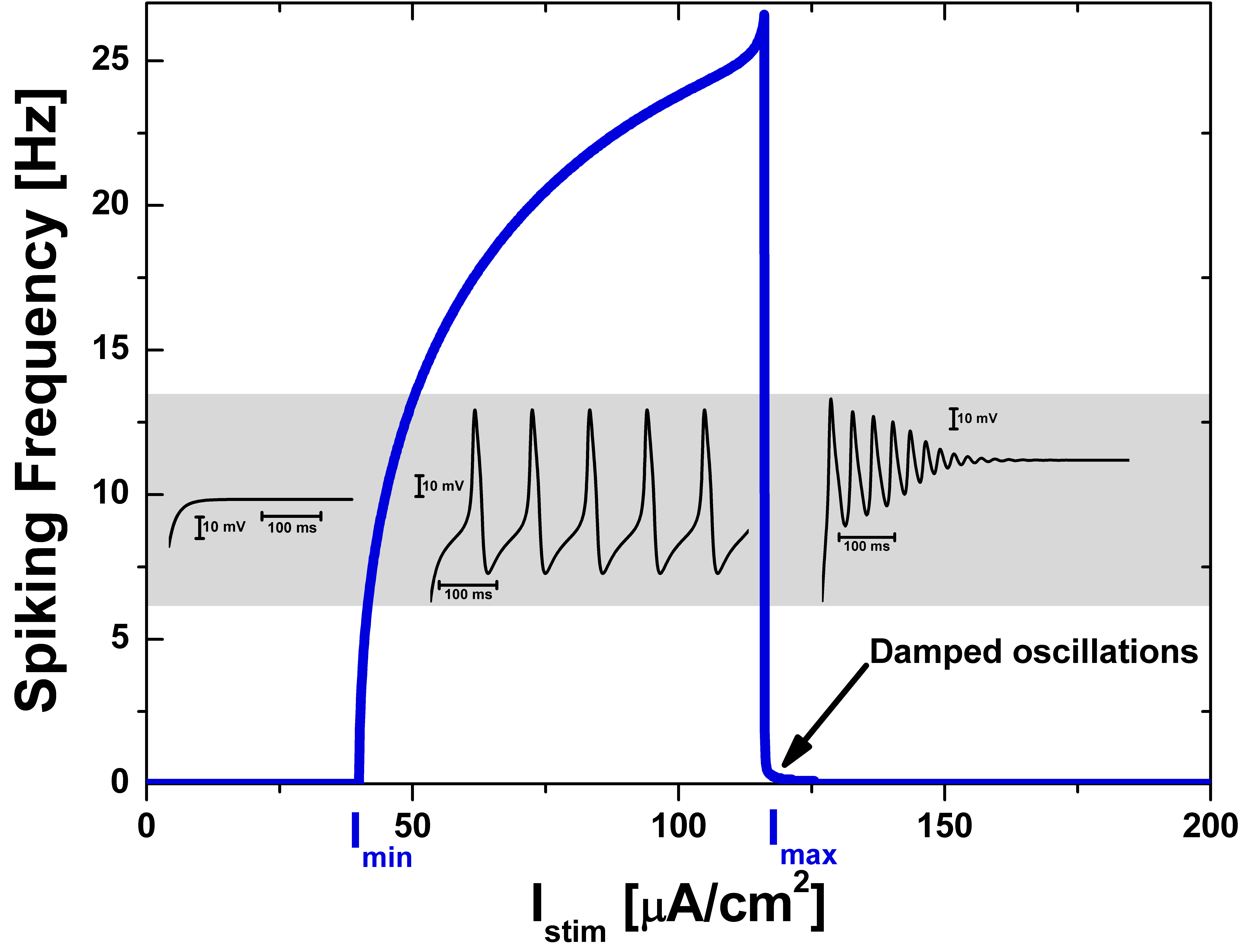

### Figure_2.png

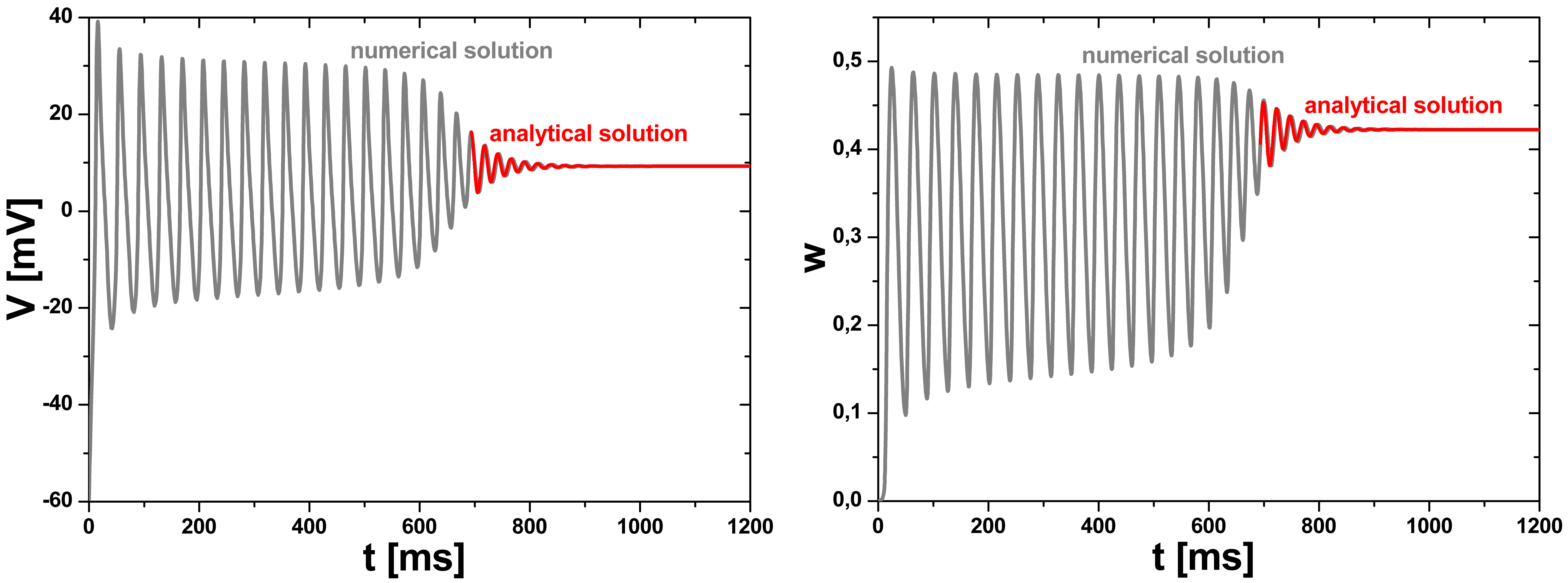

### Figure_3.png

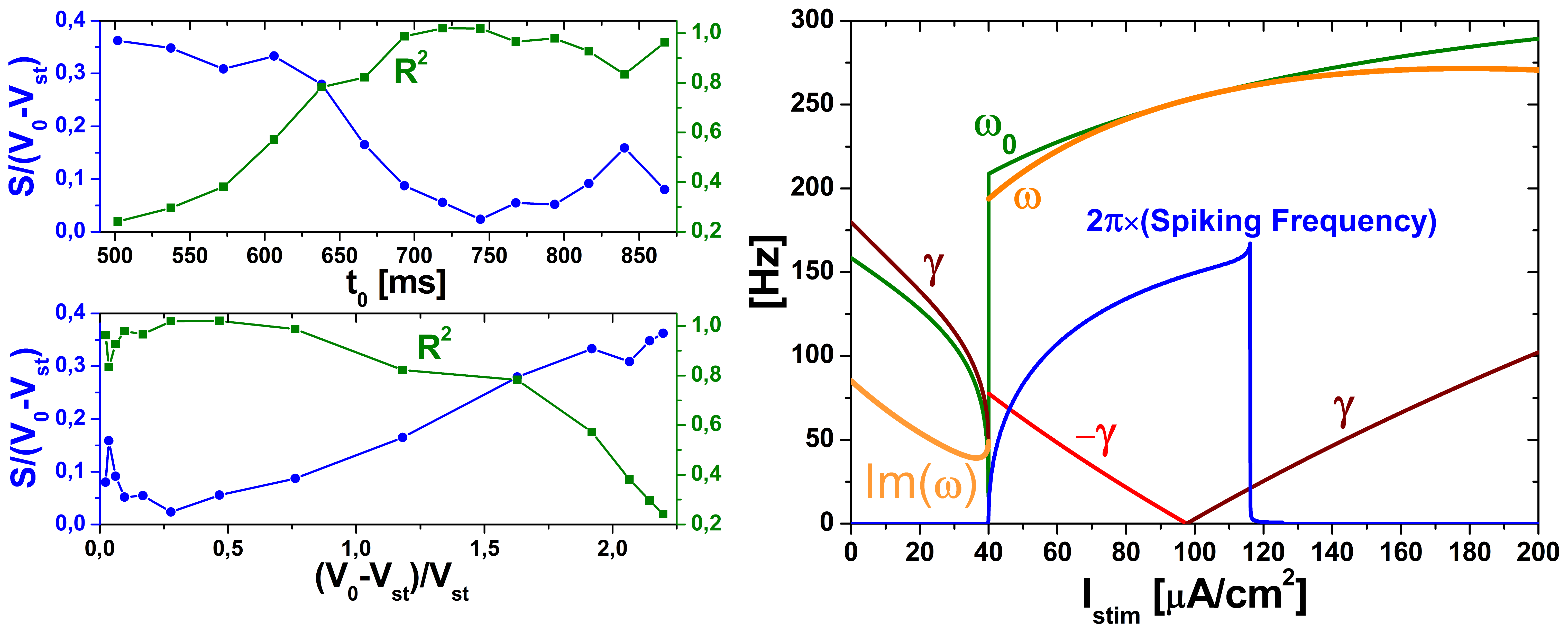

### Figure_S1_f-I_curves_ML_neuron_Exc_Types_1_and_2.png

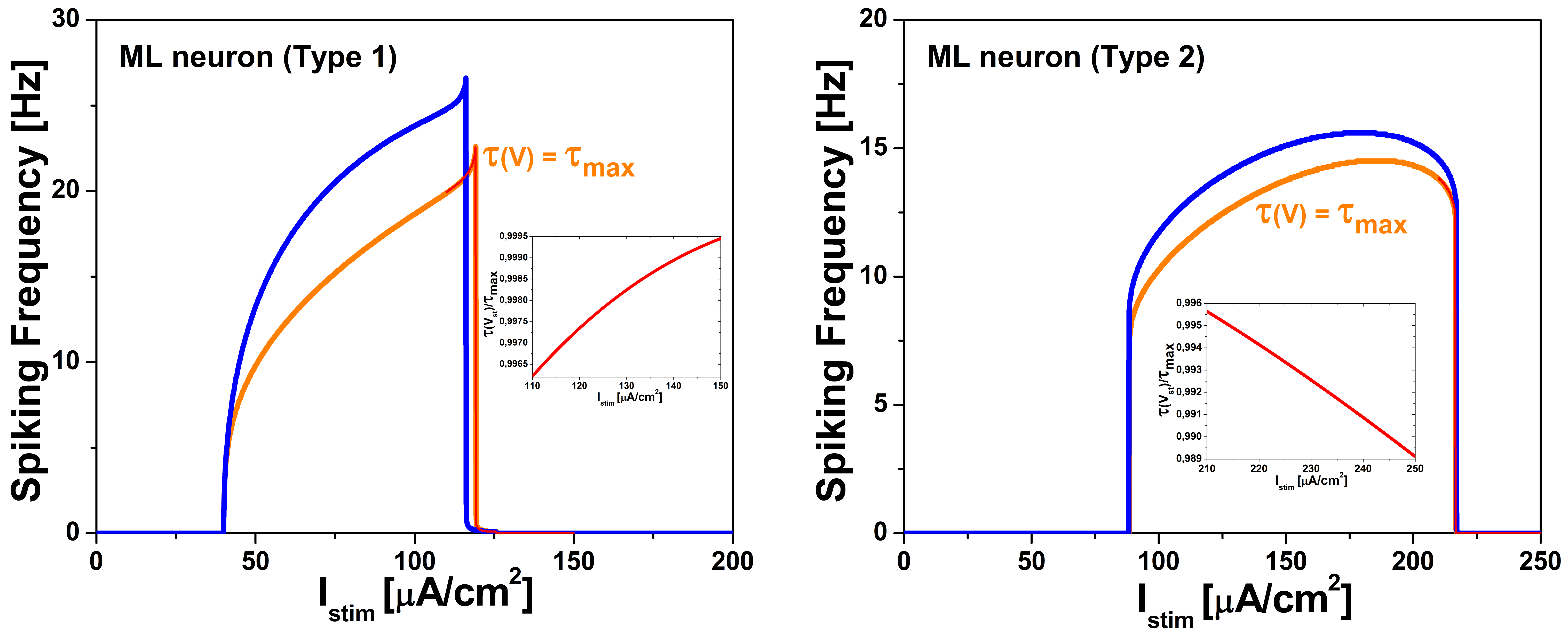

### Figure_S2_ML_neuron_dynamics_ExcType=2_FITTING_analyt_solution.png

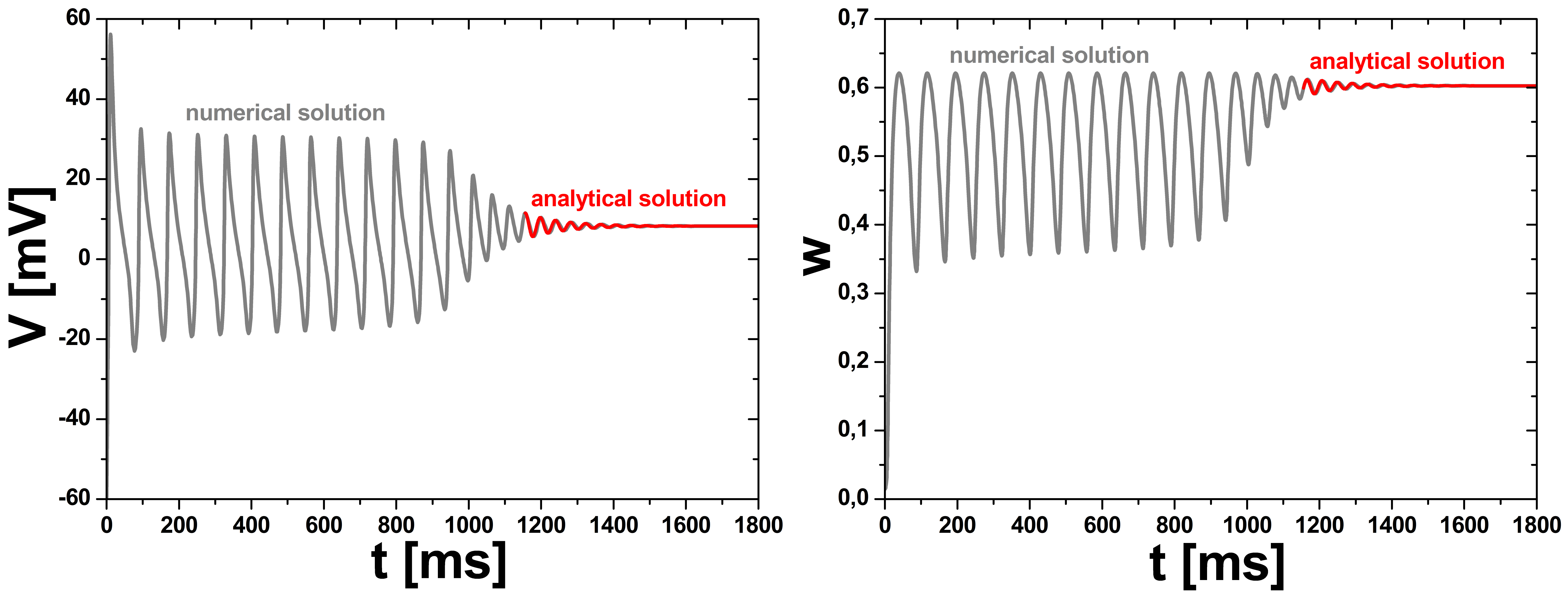
